# Minimal *N*-methylated and stapled peptide inhibitors of the autophagy protein GABARAP

**DOI:** 10.64898/2026.03.09.710535

**Authors:** Imani McDonald, Joana A. Wilms, Nicholas Cardi, Alexander Engstrom, Jiayuan Miao, Dieter Willbold, Yu-Shan Lin, R. Scott Lokey, Oliver H. Weiergräber, Joshua A. Kritzer

## Abstract

The LC3/GABARAP protein family is a promising target for selective inhibition of autophagy. Further, LC3/GABARAP ligands have been used as targeted degraders of soluble proteins, protein aggregates, mitochondria, lipid droplets, and RNA. However, the small molecules used for such applications have poor binding affinity and known off-target effects. LC3/GABARAP proteins are challenging targets for small-molecule drug development due to their long, shallow binding grooves. In this work, we evaluate multiple approaches to stabilizing the extended structure of the native binding motif, producing *N*-methylated peptides and stapled peptides with low nanomolar affinity. A crystal structure and molecular dynamics simulations support a model where the *N*-methylation pre-organizes the motif into an extended, strand-like structure. *N*-methylation allowed minimization of the binding motif to a tetrapeptide that retained sub-micromolar affinity while minimizing charge and overall molecular weight. The truncated, *N*-methylated tetrapeptide showed passive permeability in artificial membrane and cell-based transwell assays. These results highlight new drug-like space for LC3/GABARAP ligands with high affinity and subfamily selectivity.

**Table of Contents Graphic:** Drawn and developed by Mollie McGibbon and Joshua Kritzer

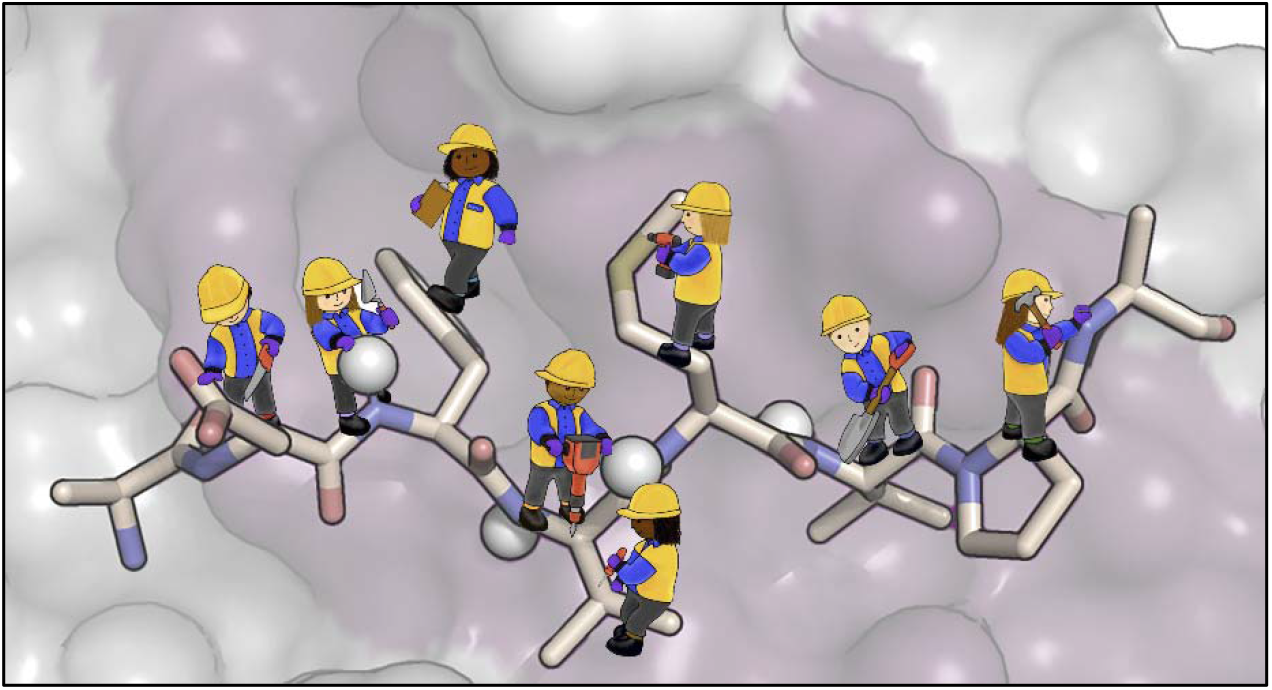

## Introduction

Macroautophagy is an essential recycling pathway in eukaryotic cells in which cytosolic material is shuttled to the lysosome for degradation.^1^ At a basal level, autophagy functions as a quality control mechanism that degrades misfolded proteins, damaged organelles, and pathogens. Under stress, such as starvation, nutrient deficiency, hypoxia, or oxidative stress, autophagy is upregulated to maintain proteostasis and to provide metabolites.^2–5^ The Atg8 protein family is critical for macroautophagy across all eukaryotes. Human Atg8 proteins include the LC3 proteins LC3A, LC3B, and LC3C and the GABARAP proteins GABARAP, GABARAPL1, and GABARAPL2/GATE16. LC3/GABARAP proteins are reversibly conjugated to membrane lipids and function as protein-protein interaction modules to mediate autophagosome initiation, elongation, closure, trafficking, and fusion with the lysosome.^6^ They are also critical for recruiting cargo to autophagosomes for degradation. Thus, LC3/GABARAP ligands have been sought both as specific autophagy inhibitors and as components of targeted protein degraders that tether proteins-of-interest to the autophagosome.^7–15^

The native protein-protein interactions of LC3/GABARAP proteins have been extensively studied.^16–19^ These interactions are almost universally mediated through a short peptide motif called the LC3-interacting region (LIR), defined as [W/F/Y]-X_1_-X_2_-[V/I/L]. Within this binding motif, specific human LC3/GABARAP paralogs have preferences for specific residues at the aromatic position, the aliphatic position, and/or at the intervening amino acids.^6,16–20^ The LIR motif binds in an extended, β-strand-like conformation, forming a short intermolecular β-sheet with LC3/GABARAP.^6,17,18,20,21^ Previous work used peptide stapling or computational design to produce peptides with high affinity for LC3B, GABARAP, or both.^22–24^ While these efforts produced peptides with high affinity, they all used larger peptide scaffolds which are unlikely to be cell-penetrant. For example, prior work from the Kritzer group produced 12-mer stapled peptides with mid-to-low nanomolar affinity, but these peptides still required micromolar concentrations to have effects on autophagy.^23^ The most potent small molecules that bind human LC3/GABARAP proteins have micromolar affinity, and most have known binding to other, non-autophagy human proteins.^25–27^

In this work, we sought to use conformational stabilization to develop smaller peptidic or peptidomimetic ligands for GABARAP that retained sub-micromolar affinity. Conformational stabilization has most commonly been investigated in α-helices using intramolecular cross-links or “staples,” and similar strategies have expanded to other structures like β-hairpins and β-sheets.^28–30^ Recently, stabilization of extended structures has been explored through intrastrand (*i,i*+2) side-chain staples by the Dawson group using dialkyne staples and by the Del Valle group using bis-thioether staples.^31,32^ Using a different approach, the Wilson group has also shown that *N*-methylation can promote binding in a β-strand conformation; detailed kinetic and thermodynamic analysis attributed the increase in binding affinity to destabilization of the unbound state.^33^ Inspired by this prior work, we sought to apply (*i,i*+2) staples and *N*-methylation to LIR peptides to develop high-affinity binders with a smaller size, with the goal of producing passively cell-penetrant compounds.

## Results

### ULK1 9-mer peptide

As a starting point, we chose a LIR peptide derived from Unc-51-like protein kinase (ULK1). ULK1is one of four proteins in the ULK1complex that initiates autophagosome formation during autophagy.^1^ The ULK1 LIR motif was attractive as a starting point because it has relatively high affinity (*K*_d_ = 50 nM for the 20-mer peptide), it has selectivity for GABARAP over LC3B, and detailed structural information was available.^20^ The crystal structure of the ULK1 LIR motif bound to GABARAP shows the characteristic extended conformation, with additional contributions from electrostatic interactions by negatively charged residues N-terminal to the LIR motif (Fig. 1).^20^ We began by truncating the larger 20-mer ULK1 peptide to 9 residues, TDDFVMVPA, which contains the canonical LIR motif FVMV (Fig. 1) and two negative charges N-terminal to the LIR motif. Prior work suggested that this 9-mer peptidewould retain most of the binding affinity for GABARAP.^20^ We substituted norleucine for methionine to avoid unwanted oxidation products, and we attached fluorescein to the N-terminus to enable fluorescence polarization binding assays as described.^22,23^ The dye-labeled 9-mer, peptide M1, had a *K*_d_ of 557 nM for binding recombinant GABARAP (Fig. 2a). We concluded that M1 would be a good starting point for testing peptide stapling and *N*-methylation strategies.

**Figure 1.**
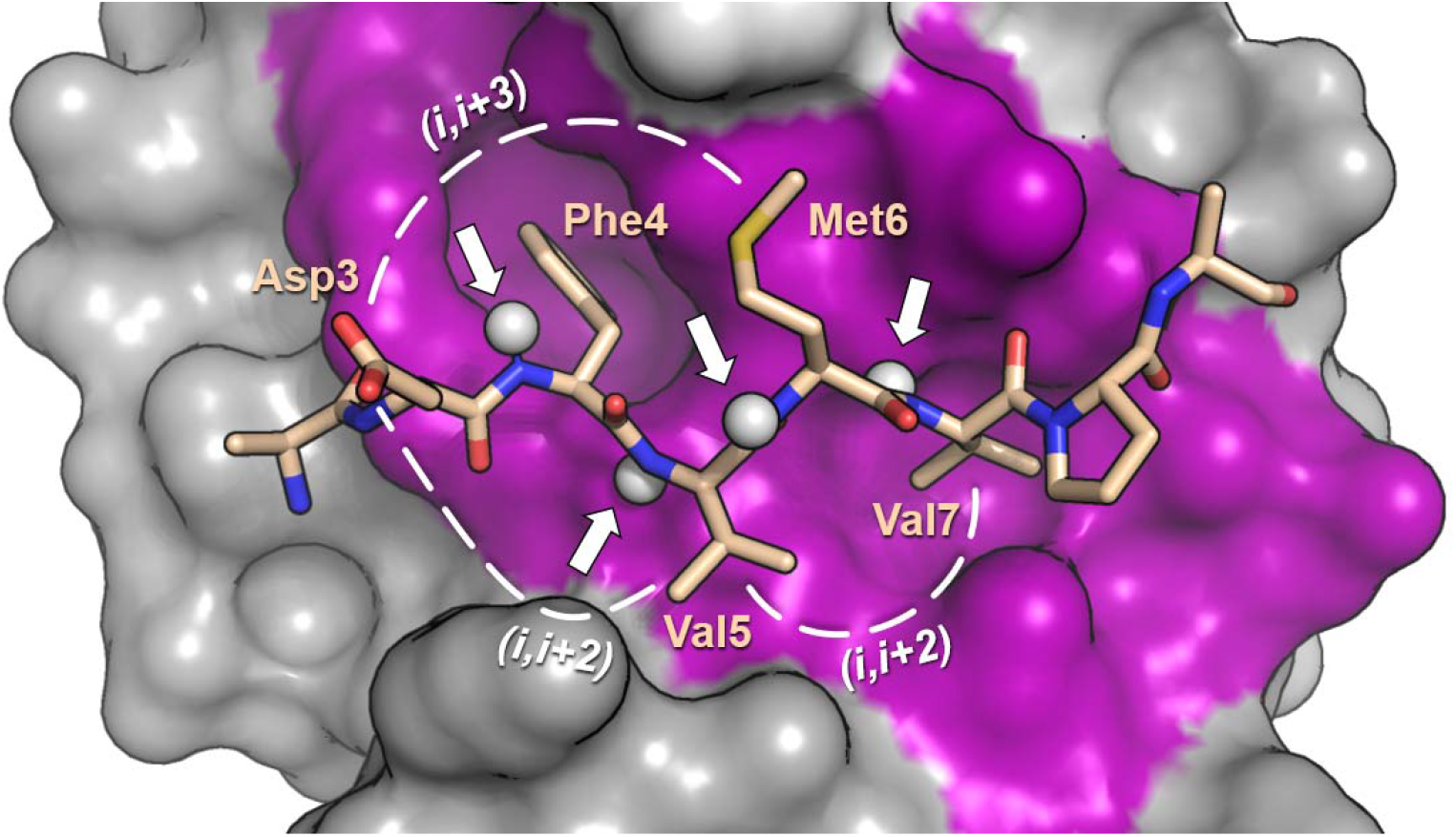
Crystal structure of a portion of ULK1 bound to GABARAP^20^. The long, shallow binding groove is highlighted in magenta. The side chain pairs Asp3/Met6, Asp3/Val5, and Val5/Val7 were judged suitable for side-chain stapling with respect to direction and distance (dotted lines). Amide protons within the core LIR motif are rendered as white spheres and indicated with arrows.

**Figure 2.**
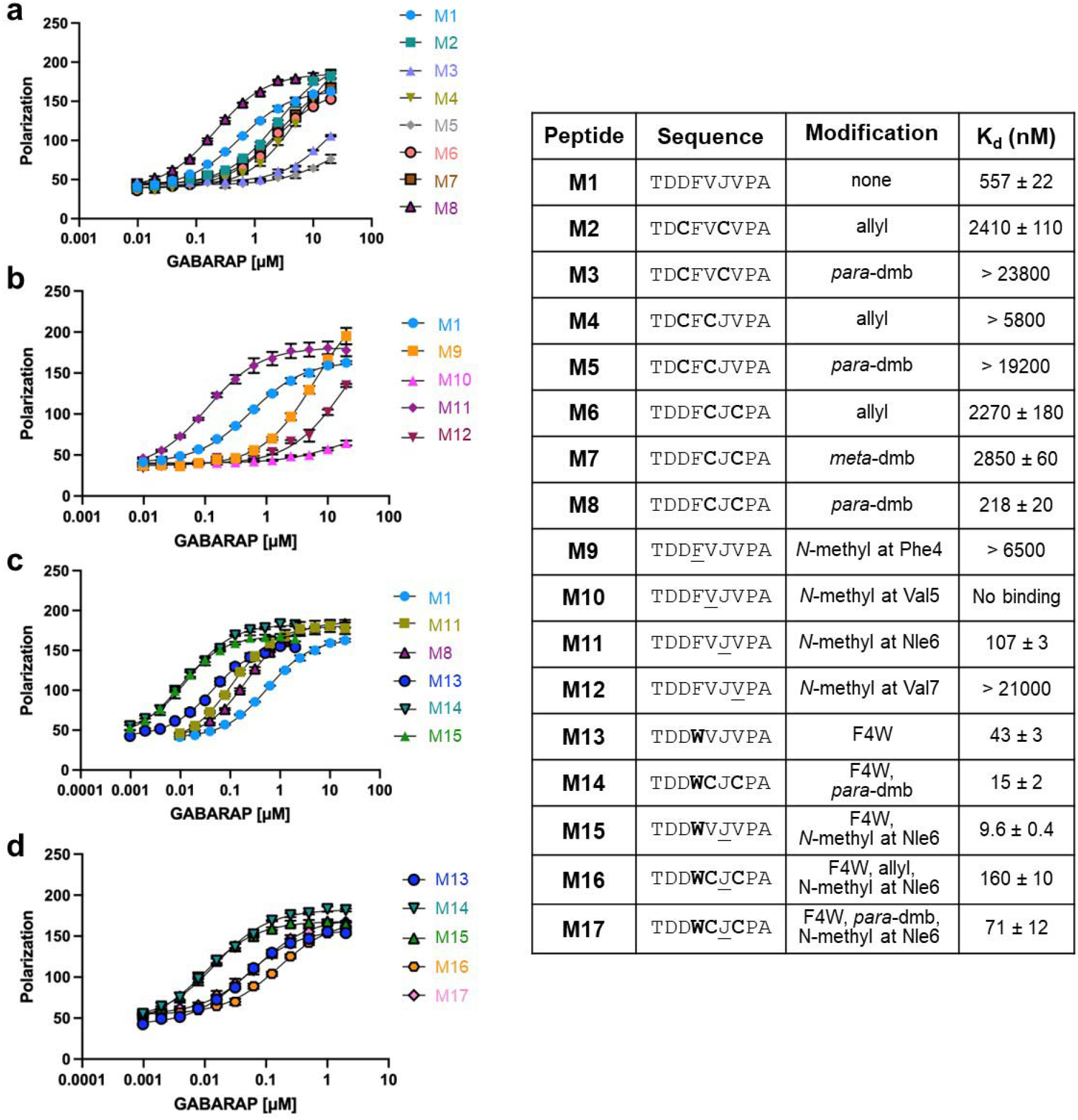
Effects of side-chain staples, *N*-methylation, and Trp substitution within the core LIR motif. **a**. Cysteine-substituted peptides were stapled with either *para*- or *meta*-dimethylbenzene (dmb), or they were allylated as an unstapled control.^34^ An (*i,i*+3) staple with *para*-dmb (M3) and an (*i,i*+2) staple with *meta*-dmb (M7) also decreased affinity. However, an (*i,i*+2) staple with *para*-dmb (M8) led to a two-fold increase in affinity. **b**. *N*-methyl residues were substituted within the core LIR motif of M1. *N*-methylation decreased affinity at three of four positions, but increased affinity five-fold at Nle6. **c**. Phe4 was substituted with Trp within the parent peptide M1 and stapled and *N*-methylated analogs. Replacement of Phe4 with Trp led to a 10-fold increase in affinity for each peptide. **d**. (*i,i*+2) stapling and *N*-methylation were combined in peptide M17. Combination of the staple and the *N*-methyl modification did not yield an additive effect on binding. Binding affinities for recombinant GABARAP were measured by fluorescence polarization using peptides labeled with fluorescein on their N-terminus. J denotes norleucine, a structural analog of methionine. Cysteines in bold are further modified as denoted in the table. The plots show averages from three independent trials and error bars show standard error of the mean.

### Stapled analogs of M1

Guided by the crystal structure of ULK1 bound to GABARAP,^20^ we designed peptides stapled at (*i,i*+2) and (*i,i*+3) positions within M1 (Fig. 1). We applied relatively rigid linkers that would promote an extended structure and preclude more compact turn structures, a strategy previously described as a “stretched” peptide.^32^ We substituted pairs of cysteine residues at several positions and stapled them with dimethylbenzene (dmb) linkers^31,34^ that roughly matched the length between the corresponding side chains in the crystal structure. We also prepared unstapled control peptides by treating the cysteine-substituted parent peptides with allyl bromide, which produced chemically similar peptides without a conformational constraint.

Using this method, we prepared cysteine-substituted, allylated control peptides M2 and M4 and stapled analogs M3 and M5. M3 was stapled at (*i,i*+3) positions, M5 was stapled at (*i,i*+2) positions, both were stapled with a *para*-dmb linkers due to the longer distance between side chains, and both were designed to avoid altering the key LIR motif residues Phe4 and Val7.^21^ The linear, allylated controls M2 and M4 each showed substantial loss of affinity compared to M1 (*K*_d_ values of 2.4 and 5.8 μM, respectively, Fig. 2a). We surmised that this result was due to the loss of one of the negatively charged Asp residues N-terminal to the LIR motif. The stapled analogs, M3 and M5, showed even poorer affinity (Fig. 2a), suggesting these substitution positions and/or staples are not compatible with binding.

We next tested a substitution position that did not remove either of the Asp residues. Substituting two cysteines at Val5 and Val7, directly within the LIR motif, we observed that allylated control M6 lost over five-fold affinity relative to the parent peptide M1 (Fig. 2a). We attributed this observation to the loss of the Val7 side chain, a key functional group within the LIR motif.^17–20^ Due to the close proximity of the two sidechains in the crystal structure, we tested analogs stapled with either *meta*-dmb or *para*-dmb. The peptide with the *meta*-dmb staple, M7, had an affinity similar to the linear, allylated control. However, the peptide with the *para*-dmb staple, M8, bound with a *K*_d_ of 218 nM, ten-fold better than its isomer M7 and about two-fold better than the parent peptide M1 (Fig. 2a). These results suggested that stapling could overcome the loss of the Val7 side chain, and that the (*i,i*+2) *para*-dmb linker was particularly compatible with GABARAP binding.

In prior work with LIR motif peptides, substitution of stapled cysteines with penicillamine, a β-branched analog of cysteine, improved binding affinities.^23^ Since stapled peptide M8 substituted Val7 in the core LIR motif with cysteine, we hypothesized that substitution of penicillamine at this position might better recapitulate native binding interactions. We substituted either or both cysteines in M8 with penicillamine and stapled them using the *para*-dmb staple. All stapled peptides with penicillamine substitutions had poorer affinity than M8 (Fig. S3). Interestingly, the allylated controls of these penicillamine-substituted peptides showed poorer affinity when either cysteine was substituted individually, indicating a preference for valine over longer β-branched residues, but slightly improved affinity when both cysteines were substituted with penicillamine (Fig. S3). These results demonstrate that, even without stapling, substitutions within the LIR motif at positions 5 and 7 have synergistic and non-additive effects. They also suggested that the *para*-dmb-stapled M8 might be binding in a conformation that is different from the parent ULK1 peptide.

### *N*-methylated analogs of M1

We next evaluated the impact of backbone *N*-methylation within the core LIR motif of M1. In the ULK1-GABARAP crystal structure, the amide protons of the core LIR valine residues (Val5 and Val7 in M1) hydrogen bond with GABARAP, forming a short intermolecular β-sheet (Fig. 1).^20^ As expected, *N*-methylation at these positions within M1 disrupted binding almost entirely (peptides M10 and M12, Fig. 2b). By contrast, the amide protons of the other core LIR residues (Phe4 and Nle6 in M1) are solvent-exposed in the ULK1-GABARAP crystal structure (Fig. 1). When Phe4 was *N*-methylated, we observed at least a ten-fold decrease in binding affinity (peptide M9, Fig. 2b). It is unclear why this substitution had such a dramatic effect. Interestingly, substitution of an *N*-methyl norleucine at position Nle6 increased binding to GABARAP five-fold, with a *K*_d_ of 107 nM (peptide M11, Fig, 2b). Because the *N*-methyl group seemed unlikely to interact with GABARAP directly, we hypothesized that the increase in affinity arose from alterations to the conformational ensemble of the unbound peptide.^33^

### Trp substitutions enhance affinity

The LIR motif contains an aromatic amino acid that binds a well-characterized hydrophobic pocket in LC3/GABARAP proteins.^6,18^ Among native LIR motifs, Phe, Tyr, and Trp are all observed in this position,^17–20^ and the native ULK1 LIR contains Phe. Previous work has found that replacing Phe or Tyr with Trp increases affinity to Atg8s by about 10-fold.^35^ Therefore, we replaced Phe4 with Trp to see if this trend would hold for M1 and its stapled and *N*-methylated analogs. Indeed, replacing Phe4 with Trp within M1 increased affinity by about 10-fold, from 557 nM to 43 nM (peptide M13, Fig. 2c). When applied to the stapled peptide, the Phe4-to-Trp substitution also improved affinity by about ten-fold, improving affinity to 15 nM (peptide M14, Fig. 2c). Similarly, when applied to the *N*-methylated peptide, the Phe4-to-Trp substitution improved affinity from 107 nM to 9.6 nM (peptide M15, Fig. 2c). Thus, we found that the Trp substitution at position 4 improved affinity in all contexts and we retained it in continued designs.

### Stapling and *N*-methylation are not compatible

After finding that *N*-methylation and (*i,i*+2) stapling with *para*-dimethylbenzene individually increased affinity to GABARAP, we asked if these effects are additive. Combining these modifications yielded a peptide with a *K*_d_ of 71 nM for GABARAP, which was poorer than either modification on its own (peptide M17, Fig. 2d). The allylated analog, peptide M16, had an even poorer binding affinity, demonstrating that the staple was still beneficial to binding. However, it was striking that the staple improved binding 10-fold in peptides that lacked *N*-methylation (M8 compared to M6, Fig. 2a), but the staple only improved binding 2-fold in peptides with an *N*-methyl Nle6 (M17 compared to M16, Fig. 2d). These results suggested that *N*-methylated analog M11 and stapled analog M8 may have different binding modes and/or different solution ensembles that affect free energy of binding.

To explore these possibilities, we performed molecular dynamics simulations on minimal peptides that represented the core LIR domains of M1, stapled M8, *N*-methylated M11, and stapled and *N*-methylated M17. We retained the original Phe4 residue for consistency. Analysis of the backbone (□, *ψ*) dihedral angles revealed that (*i,i*+2) stapling and *N*-methylation each promote an extended structure with dihedrals largely in the β-sheet and PPII regions, close to the bound conformation (Fig. 3). This finding is consistent with the observed enhancement in binding affinity of each modification on its own. However, large differences are observed for the dihedral angles of the central norleucine residue. When stapled, the norleucine shifts slightly, preferring extended structures with slightly larger negative □ values (Fig. 3, second row, Nle residue). When *N*-methylated, the norleucine shifts more dramatically, favoring positive □ values (Fig. 3, third row, Nle residue). When the peptide is both stapled and *N*-methylated, the norleucine prefers dihedral angles in a region between the β/PPII and α-helix regions, different than the preferred regions for the stapled peptide or the *N*-methylated peptide (Fig. 3, fourth row, Nle residue). These results suggest that *N*-methylation and stapling preorganize the backbone into different conformational ensembles, and that combining the two modifications promotes a conformational ensemble that is distinct from the ensembles of peptides with individual modifications. We noted that the results for the *N*-methylated peptide are somewhat counterintuitive, as *N*-methylated L-amino acids are generally expected to prefer negative □ values.^36,37^ These results are also inconsistent with the co-crystal structure of the *N*-methylated peptide bound to GABARAP, where the □ value of the *N*-methylated norleucine is –86° (purple lines in Fig. 3). To test whether these results were due to a force field artifact, we examined the conformational ensemble of an *N*-methylated alanine dipeptide using several force fields: RSFF2,^38^ Amber14SB,^39^ Amber19SB,^40^ OPLS2005,^41^ and OpenFF.^42^ In all simulations, the □ angles predominantly favored positive values. Future work is required to improve modeling for structural ensembles of *N*-methylated peptides. Still, these results provided further support for a model in which the *N*-methylated peptide M11 and the stapled peptide M8 bind GABARAP in different conformations.

**Figure 3.**
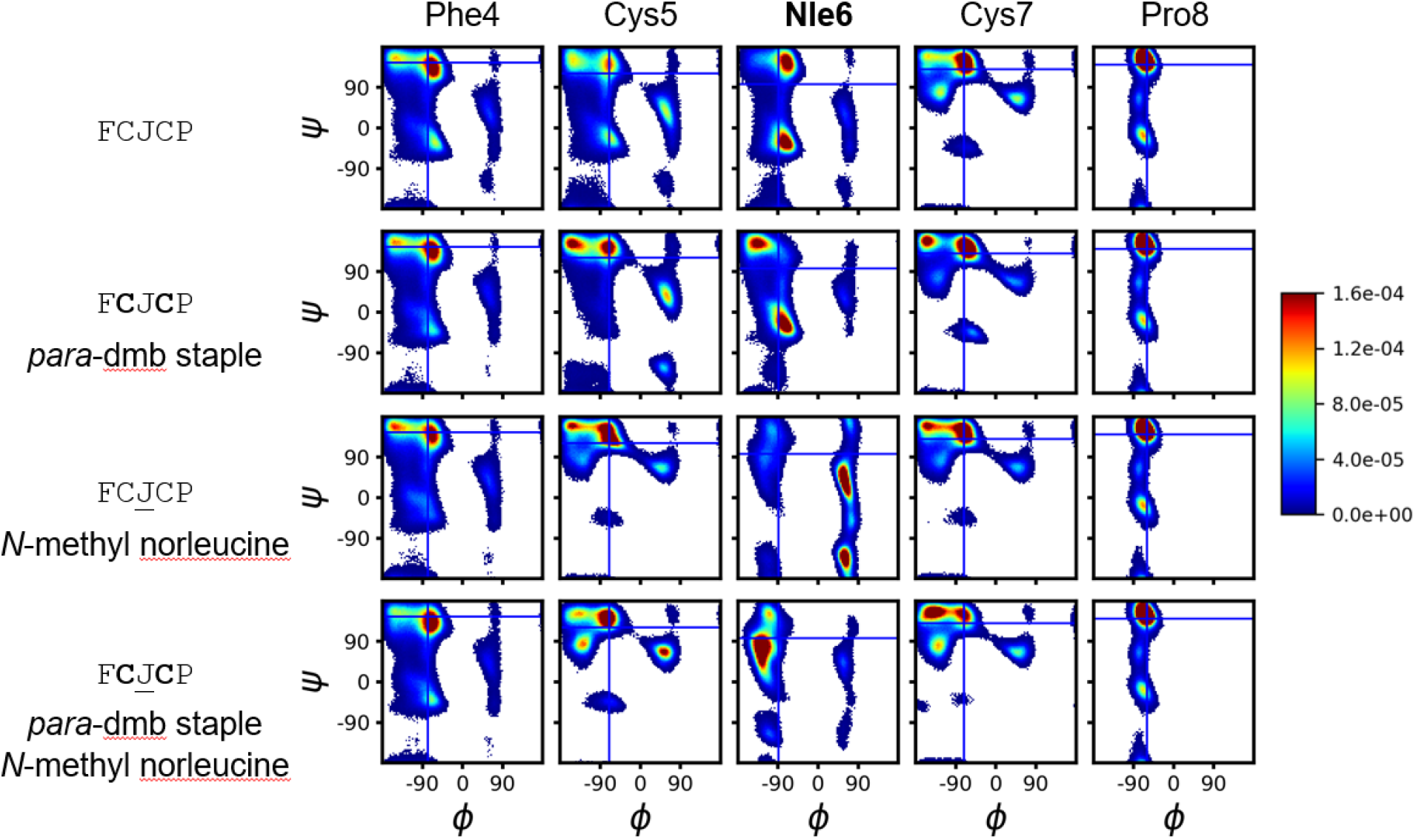
Molecular dynamics (MD) simulations of model peptides. All-atom, explicit-solvent molecular dynamics of model pentapeptides were performed using bias-exchange metadynamics to ensure exhaustive sampling. Global ensembles were simulated for peptides with no modification, a *para*-dimethylbenzene (dmb) staple, an *N*-methyl norleucine (<u>J</u>), or both modifications. Peptides had acetylated N-termini and amidated C-termini. Residues are numbered in correspondence with their positions in M1 and analogs. Ensembles are shown as Ramachandran density plots for each amino acid. For comparison, the bound conformation of the corresponding residue of the ULK1 peptide (Fig. 1) is indicated with blue lines.

### Crystal Structure of M15 bound to GABARAP

We were interested in the binding modes of stapled peptide M14 (*K*_d_ = 15 nM) and *N*-methylated peptide M15 (*K*_d_ = 9.6 nM), so we pursued X-ray crystallography. Only M15 yielded usable crystals, and we acquired a crystal structure of M15 bound to GABARAP with 1.42 Å resolution. The crystal structure revealed that M15 binds in an almost identical mode to the parent ULK1 peptide (Fig. 4),^20^ with a backbone heavy-atom RMSD of 0.1 Å for the core LIR motif. The *N*-methyl at Nle6 points out towards solvent, and Trp4 is positioned similarly to the corresponding Phe in the ULK1 peptide. Slight differences were observed in the positions of the flanking residues Ala9, Asp2, and Asp3, but it is unclear whether that was due to the *N*-methylation and Trp substitution, or whether it was due to crystallization conditions.

**Figure 4.**
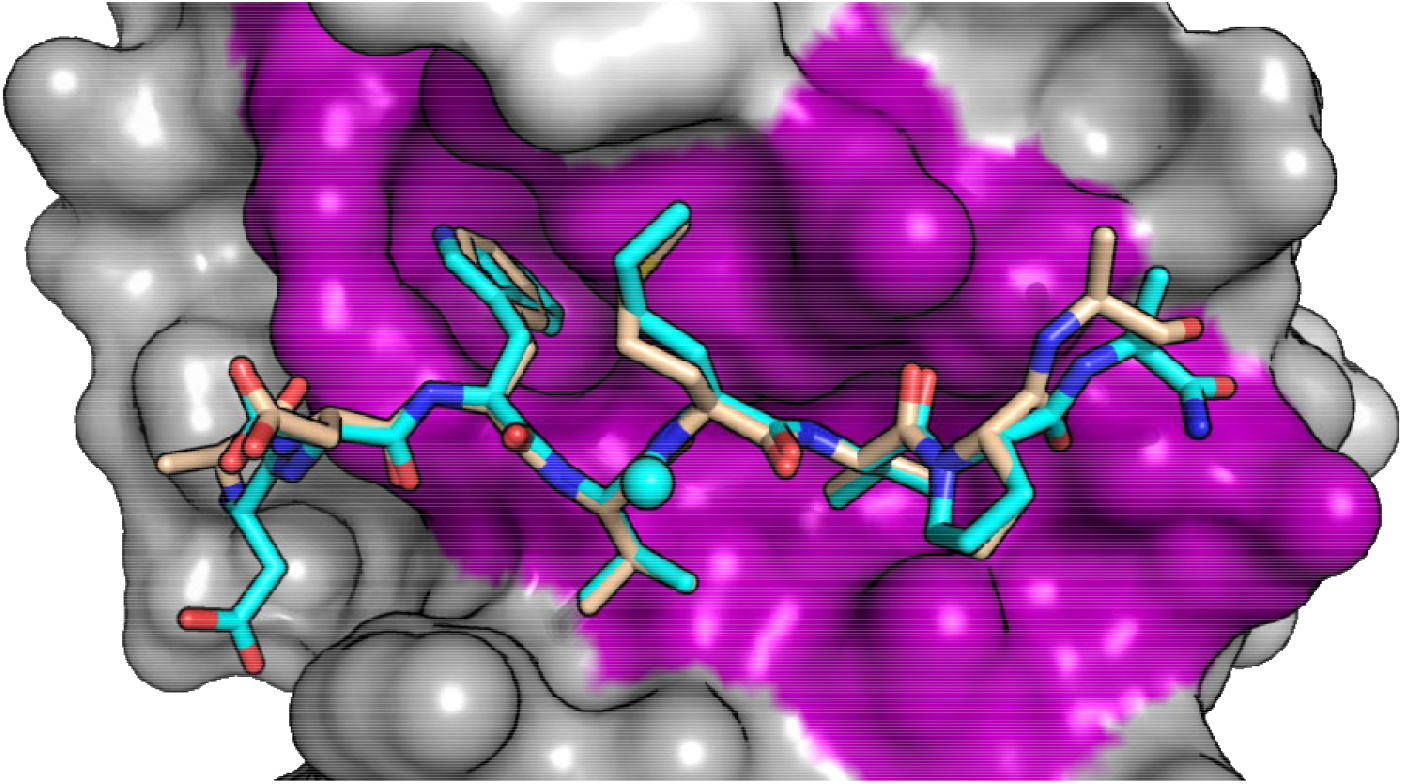
Crystal structure of *N*-methylated peptide M15 bound to GABARAP. GABARAP-bound M15 (cyan) is positioned similarly to the corresponding portion of GABARAP-bound ULK1 peptide (wheat).^20^ The *N*-methyl at Nle6 points towards solvent (cyan sphere) and Trp4 engages the expected hydrophobic pocket. The core LIR motifs overlay with a tight backbone RMSD of 0.1 Å, but slight variations from the ULK1 conformation are observed in the positioning of the N- and C-termini of M15. The final coordinates have been uploaded to the Protein Data Bank with ID 9I9X, with M15 denoted as peptide IM-2.

### Truncation of stapled and *N*-methylated peptides

While *N*-methylation and stapling should offer some resistance to peptidases, they are not sufficient to promote passive cell penetration.^44^ These 9-mer peptides still have a high molecular weight, a high number of hydrogen bond donors, and two negative charges, making it extremely unlikely they will penetrate cell membranes. Because we observed low nanomolar affinity for our *N*-methylated and stapled peptides, we sought to truncate them to maintain sub-micromolar affinity while steering towards more passively permeable chemical space. Truncation of the N-terminus of the stapled peptide M14 led to a roughly 5-fold loss in affinity, which was expected due to the loss of both Asp residues (peptide M18, Fig. 5a). Perhaps more surprisingly, truncating the C-terminal Pro-Ala residues led to a similar loss in affinity (peptide M19, Fig. 5a), despite the relative lack of importance of these residues for canonical LIR binding. Combining these truncations left only the core LIR sequence, WCJC, with the cysteines stapled with *para*-dimethylbenzene. This stapled tetrapeptide, M20, had even poorer affinity than either truncation alone (greater than 1.6 µM).

**Figure 5.**
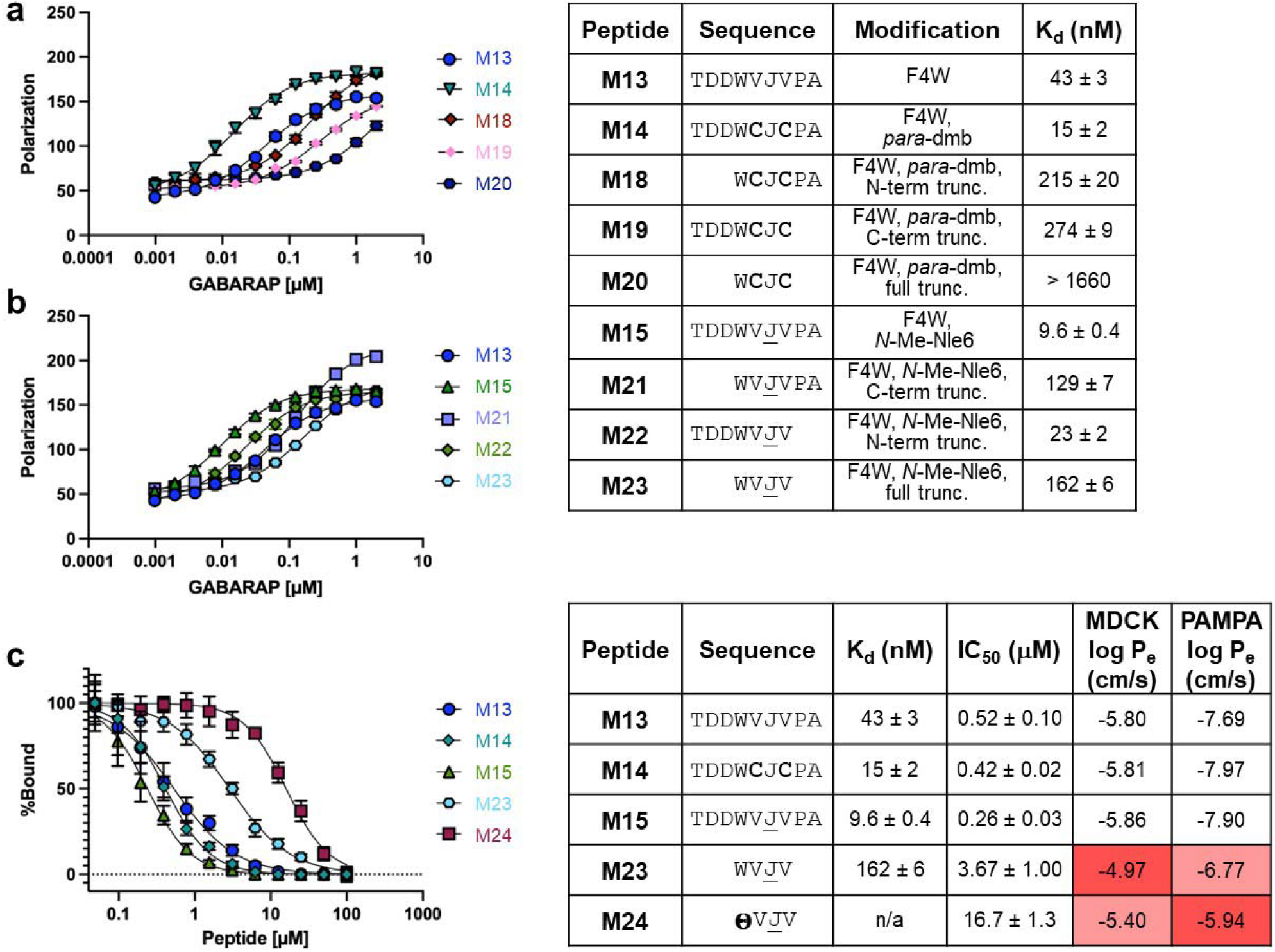
Effects of truncation on stapled LIR peptides and *N*-methylated LIR peptides. **a**. Truncated analogs of stapled peptide M14. Truncation at either of the termini led to a ~15-fold decrease in affinity. Simultaneous truncation at both termini led to an even larger loss in affinity. **b**. Truncated analogs of *N*-methylated peptide M15. Truncation at the N-terminus led to a 13-fold decrease in affinity, but truncation at the C-terminus led to only a 2-fold decrease in affinity. Simultaneous truncation of both termini led to a roughly 16-fold loss in affinity. **c**. Non-dye-labeled peptides were tested for inhibitory potency and for passive penetration using PAMPA and MDCK assays. The inhibition assay was an AlphaScreen assay which measured inhibition of the interaction between GABARAP and a high-affinity peptide ligand, K1.^23,43^ **θ** represents an indole 3-acetic acid cap, similar to Trp but without the N-terminal amino/amide group. J denotes norleucine, *N*-methylated residues are underlined, and cysteines in bold are further modified as denoted in the table. The plots show averages from three independent trials and error bars show standard error of the mean.

Truncation of the *N*-methylated peptide M15 showed different results. Truncation of the three N-terminal residues resulted in a 13-fold loss in affinity (peptide M21, Fig. 5b) while truncation of the two C-terminal residues resulted in only a 2-fold loss in affinity (peptide M22, Fig. 5b). These results were more consistent with prior work on ULK1-derived LIR peptides and other LIR peptides,^17–20^ and the different effects of truncation on the stapled peptides compared to the *N*-methylated peptides further support a model where these peptides bind GABARAP in different conformations. Combining the two truncations produced peptide M23, an *N*-methylated tetrapeptide containing only the core LIR motif. M23 had a *K*_d_ of 162 nM, about 15-fold higher than full-length *N*-methylated peptide M15. While this was a significant loss in binding, it was promising that the peptide retained sub-micromolar affinity when truncated to such a small length, and without negatively charged residues.

Our final design iteration sought to remove the N-terminal amide of the *N*-methylated tetrapeptide. We surmised that this amide might not be needed for GABARAP binding, and that removal of the amide could further increase passive cell permeability. Because we had been using the N-terminus to attach fluorescein for fluorescence polarization assays, we used an AlphaScreen competitive binding assay^45^ to test variants lacking an N-terminal amide. The AlhpaScreen assay measures the relative potency for compounds to inhibit the binding between GABARAP and K1, a high-affinity ligand discovered via phage display.^23,43^ We prepared full-length peptides with the Phe4-to-Trp substitution (M13), the (*i,i*+2) *para*-dimethylbenzene staple (M14), and an *N*-methylated Nle6 (M15) as comparators. We also prepared the truncated *N*-methylated tetrapeptide (M23) and a variant in which the N-terminal acetylate-tryptophan was replaced by an 3-indole-acetic acid (M24), which effectively removes the N-terminal amide of the tetrapeptide. Full-length peptides M13, M14, and M15 had IC_50_ values of 519, 418, and 256 nM, respectively (Fig. 5c). These IC_50_ values were roughly 10-to 20-fold higher than their *K*_d_ values, which was consistent with prior results with this assay.^45^ The *N*-methylated tetrapeptide M32 had an IC_50_ value of 3.67 μM, also 20-fold higher than its *K*_d_. Finally, we tested the analog lacking the N-terminal amide (M24), which had an IC_50_ value of 16.7 μM, roughly five-fold worse than M23. This affinity loss suggests that either the Trp’s N-terminal amide is involved in subtle contacts with GABARAP, or that the chirality at the Trp helps pre-organize the side chain for GABARAP binding. Either way, we observed a tradeoff between further truncation of the backbone and binding affinity at the N-terminus of the tetrapeptide.

### Selectivity for GABARAP over LC3B

The ULK1 peptide is somewhat unique among LIR-motif peptides for being highly selective for GABARAP.^17,18,20^ We sought to determine whether the stapled, *N*-methylated, Trp-substituted, and truncated peptides retained this selectivity by testing them in fluorescence polarization assays with recombinant LC3B.^22,23^ Most peptides showed no binding or minimal binding to recombinant LC3B up to 2 μM protein (Fig. S4, Table S5). These results showed that, throughout all modifications, the ULK1-derived peptides maintained at least 200-fold selectivity for GABARAP over LC3B.

### Passive permeability

GABARAP ligands M23 and M24 have molecular weights below 575 Da and they have only four and three backbone amides, respectively. Their relatively small size and low number of hydrogen bond donors suggested they might be passively cell-permeable.^46,47^ To measure passive permeability, we used two common transwell permeability assays: the parallel artificial membrane permeability assay (PAMPA) and a Madin-Darby canine kidney (MDCK) cell monolayer permeability assay.^48^ Testing the 9-mer peptide M13, the 9-mer stapled peptide M14, and the 9-mer *N*-methylated peptide M15, we observed negligible passive permeability in PAMPA and a moderate amount of permeability in the MDCK assay, perhaps due to active transport. By contrast, the tetrapeptide M23 and the 3-indole-acetic-acid-capped tripeptide M24 each showed moderate membrane permeability in PAMPA and enhanced permeability in the MDCK assay (Fig. 5c). Interestingly, M24 showed less permeability than M23 in the MDCK assay, but greater permeability in the PAMPA assay. Overall, these data demonstrated that truncation produced *N*-methylated LIR peptides with submicromolar inhibitory potency and moderate passive permeability.

## Discussion

Conformational stabilization of peptides has long been used to produce potent peptidomimetics that block protein-protein interactions.^33,49^ While much prior work has focused on peptides that bind in α-helical conformations, a growing body of work has investigated conformational stabilization of β-strands, polyproline helices, and other extended structures.^32,50,51^ In this work, we extended this effort to LIR motifs, producing nanomolar binders of GABARAP using both stapling and backbone *N*-methylation strategies.

We found that *N*-methylation did not alter bound structure, yet it increased binding affinity by 5-fold. While *N*-methylation is a common peptide modification, most prior work involved *N*-methylation of cyclic peptides, helical peptides, turn regions, or heterochiral backbones.^49,52–55^ Recent work by Wilson and coworkers has provided the most comprehensive analysis to date of the effects of *N*-methylation on the thermodynamics of β-strand preorganization and binding.^33^ This work suggested that linear *N*-methylated peptides have a lower conformational entropy in the unbound state, leading to entropy-driven improvements in binding kinetics (on-rates) and thermodynamics (free energies of binding). Our results, supported by MD simulations, support a model in which *N*-methylation similarly decreases the conformational entropy of unbound LIR peptides, leading to improved free energy of binding.

Recent work by del Valle and coworkers using a model β-hairpin peptide suggested that, while (*i,i*+2) stapling with *para*-dmb is compatible with β-strand conformation, *meta*-dmb is more β-strand-stabilizing.^31^ However, for GABARAP binding, we observed that the peptide with the *meta-*dmb staple lost affinity relative to an unstapled control, while the peptide with the *para*-dmb staple gained affinity. This effect may be due to interactions between the staple and GABARAP, as observed for much larger stapled peptides that were previously reported.^23^ We were unable to produce a crystal structure to evaluate this hypothesis directly. Alternatively, the preference for the *para*-dmb staple could be due to the slightly distorted β-strand conformation predicted by MD simulations at the Nle residue (Fig. 3). Several comparisons to the *N*-methylated peptides further support the likelihood that the stapled peptides have an altered binding conformation, including the incompatibility of the (*i,i*+2) staple with *N*-methylation at Nle6, the different structural ensembles predicted by the MD simulations, and the surprising decrease in affinity when the stapled peptide was C-terminally truncated (peptide M19 compared to M14). Further, we have demonstrated in previous work that the GABARAP binding site can undergo induced fit to allow for non-canonical binding modes.^23,43^ All together, these observations provide multiple independent lines of evidence that the stapled peptides bind GABARAP in a different conformation than the canonical, β-strand LIR motif.

In this work, we minimized the ULK1 peptide into a passively penetrant, GABARAP-specific ligand with low molecular weight, no charged residues, and as few as three backbone hydrogen bond donors. However, there is still room for further improvement. The methionine side chain adopts two conformations in the original crystal structure^29^ and the same is true for its *N*-methyl norleucine analog in the M15 complex (this study), suggesting that a larger side chain in this position might improve affinity. Additionally, further functionalization of the indole was not explored, but prior work on indole ligands suggests that non-canonical Trp analogs might improve affinity.^56^ Finally, successful minimization of the ligand to a tetrapeptide will facilitate further development using solution-phase synthesis, allowing more extensive modifications to the backbone including substitutions of amide bond isosteres at the internal amide bonds and appending more varied functional groups to the termini.^57,58^

In multiple recent studies of targeted protein degradation, tethering proteins-of-interest to GABARAP resulted in particularly efficient autophagic degradation.^59,60^ Small molecule ligands of GABARAP continue to be of high interest both as autophagy modulators and as components of targeted degraders. However, this is an especially challenging protein-protein interaction due to its shallow groove, and existing small molecule compounds generally have micromolar affinity and known off-target interactions.^25,26,45^ We expect continued development of peptidomimetic GABARAP ligands will furnish compounds with higher affinity, better cell penetration, and fewer off-target effects than current compounds. Fully optimized compounds will be useful for exploring autophagy modulation in disease models, and for developing targeted protein degraders with greater potency and drug-likeness.

## Supporting information

Complete materials and methods, peptide characterization, raw data and curve fits for all assays (PDF).

## Acknowledgments

The authors would like to thank the staff of the ESRF and EMBL Grenoble for assistance and support in using beamline ID30A-3 under proposal number MX-2587.

