## supporting information for "Minimal *N*-methylated and stapled peptide inhibitors of the autophagy protein GABARAP"

### Materials and Methods

#### Peptide synthesis and purification

Peptides were synthesized by standard Fmoc solid phase peptide synthesis using either an automated Tribute peptide synthesizer (Gyros Peptide Technologies) or an automated Prelude peptide synthesizer (Gyros Peptide Technologies). Peptides were synthesized on Rink amide resin. Fmoc deprotection was done using 20% piperidine in DMF. Coupling steps used 4 equivalents of amino acid, 4 equivalents of coupling reagent (HBTU or HATU and HOBt or HOAt), and 8 equivalents of *N,N*-diisopropylethylamine (DIPEA). After synthesis, the resin was washed with DMF, DCM, and MeOH, then dried over vacuum. All peptides were globally deprotected and cleaved from the resin with a TFA cleavage cocktail (TFA/$H_2O$/EDT/TIPS 95:2:2:1) for 3 h. After cleavage, peptides were precipitated in cold diethyl ether and centrifuged yielding a peptide pellet. The pellet was dried with nitrogen, redissolved in ACN:$H_2O$ (50:50), and lyophilized.

After lyophilization, the peptides were resuspended in ACN:$H_2O$ (50:50) for reverse-phase HPLC purification. Peptides were purified using either a C8 or C18 column with a gradient of 5 to 100% acetonitrile with 0.1% TFA over 20 min. After purification, peptides were analyzed via analytical HPLC. All peptides were used at a minimum of 95% purity as measured by analytical HPLC. Peptide mass was analyzed using MALDI-TOF mass spectrometry with 20 mg/mL $\alpha$-cyano-4-hydroxycinnamic acid in ACN:$H_2O$ (70:30) with 0.1% TFA as the matrix.

Fluorescein-tagged peptides were prepared with the sequences shown, followed by two β-alanine residues coupled to the N-terminus to serve as a linker. After these couplings, the peptide was deprotected and a coupling with 3 equivalents of 5/6-carboxyfluorescein succinimidyl ester (NHS-fluorescein, Thermo Fisher) was performed with 6 equivalents DIPEA in DMF for 1 h. The resin was washed with DMF and a second coupling under the same conditions was done overnight. After the second coupling, the resin was washed and dried, and the peptide was cleaved, lyophilized, and purified as described above.

Acetylated peptides were prepared with the sequences shown, followed by a coupling with 50 equivalents acetic anhydride and 100 equivalents DIPEA in DMF for 20 min. After coupling, the resin was rinsed and dried, and the peptide was cleaved, lyophilized, and purified as described above.

For stapling reactions, lyophilized peptide was dissolved in 50% acetonitrile, 50% aqueous buffered ammonium carbonate, pH 8. 1.1 equivalents of dibromomethylbenzene or 2.2 equivalents of allyl bromide were added to the peptide. After three hours, the reaction was quenched with 0.2% TFA and the peptide was lyophilized and purified as described above.

**Protein expression and purification**

For binding assays, recombinant His-tagged GABARAP and LC3B were expressed in BL21 (DE3) *E. coli* cells as described.[1–3] The bacteria were transformed with pET-15b expression plasmids encoding each protein. Transformed cells were plated on ampicillin agar plates and incubated at 37 ºC overnight. Individual colonies were picked and grown in 5 mL LB culture medium with 1% ampicillin at 37 ºC, shaking overnight. The 5 mL culture was then added to 1 L of LB medium with 1% ampicillin and incubated with shaking at 37 ºC until the $OD_{600}$ measured ≥ 0.6. Protein expression was induced with 1 mL of 0.5 mM Isopropyl β-D-1-thiogalactopyranoside (IPTG). Cultures were incubated for 3 hours with shaking at 37 ºC. Cells were then pelleted and stored at -80 ºC.

For protein purification, frozen cell pellets were resuspended in lysis buffer (25 mM HEPES, 150 nM NaCl, 10 mM imidazole, 0.2% lysozyme, 1 protease inhibitor cocktail pellet, and 0.001% universal nuclease). The resuspended cells were sonicated, then centrifuged to separate the lysate from the cellular debris. The clarified lysate was purified via batch affinity purification with HisPur Ni-NTA resin (Thermo Fisher Scientific). The resin was incubated with the lysate at 4 ºC for one hour, then washed with 25 mM HEPES, 500 mM NaCl, and 20 mM imidazole. After, it was washed with lysis buffer to reduce the salt from 500 mM NaCl to 150 mM NaCl. The protein was eluted from the resin with elution buffer (25 mM HEPES, 150 mM NaCl, and 500 mM imidazole). To remove imidazole, buffer exchange was performed via a desalting column into storage buffer (25 mM HEPES, 150 mM NaCl, pH 7.3). Protein purity and mass were assessed using SDS-PAGE. If needed, proteins were further purified by size exclusion chromatography using storage buffer. Protein was aliquoted and flash frozen in liquid $N_2$, then stored at -80ºC.

For crystallization experiments, GABARAP was expressed as glutathione *S*-transferase (GST) fusion protein after transforming *E. coli* BL21(DE3) T1 cells with pGEX4T2-GABARAP plasmids. Bacteria were cultivated in LB medium containing 100 µg/mL ampicillin at 37 °C until an $OD_{600}$ of 0.6–0.8 was reached. Then gene expression was induced by adding 1 mM IPTG; cultures were transferred to 25 °C and protein expression could proceed for 20 h. Afterward, cells were harvested by centrifugation at 3,000×*g* for 30 min at 4 °C. The bacterial pellet was washed with PBS (137 mM NaCl, 2.7 mM KCl, 1.8 mM $KH_2PO_4$ and 10 mM $Na_2HPO_4$) and resuspended in lysis buffer (PBS supplemented with 5% (v/v) glycerol, 0.01% (v/v) β-mercaptoethanol, 10 µg/mL DNase (AppliChem, A3778) and cOmplete EDTA-free protease inhibitor cocktail (Roche, 11836170001)) before application to the cell disruptor (Constant Systems, model TS1.1) for three cycles with 1.9 kbar at 4 °C. Lysates were cleared by centrifugation at 4 °C with 45,000×*g* for 45 min. The GST fusion protein was purified from the supernatant by affinity chromatography using Glutathione Sepharose 4B (Cytiva, 1705605). Cleavage with thrombin (Sigma-Aldrich, 1.12374) during dialysis against 10 mM Tris-HCl and 150 mM NaCl (pH 7.0) at 4 °C overnight yielded a 119-amino-acid protein carrying an N-terminal Gly-Ser extension in addition to the native sequence of the GABARAP protein. The digested and dialyzed GABARAP sample was applied to a Hiload 26/60 Superdex 75 preparatory-grade size-exclusion column (GE Healthcare) equilibrated with 10 mM Tris-HCl and 150 mM NaCl (pH 7.0) for removal of the cleaved GST-tag and further purification. Protein purity was assessed by SDS-PAGE and Coomassie staining. Fractions containing the eluted GABARAP protein were concentrated to 3–5 mg/mL using Vivaspin 20 concentrators with a 3 kDa cutoff (Sartorius), flash-frozen in liquid $N_2$ and kept at −80 °C until usage.

#### Fluorescence polarization assay

Fluorescein-labeled peptides were diluted in assay buffer (25 mM HEPES, 150 mM NaCl, 1 mM EDTA, and 0.1% Tween-20, pH 7.3) to 10 nM and 10 μL was added in triplicate to a black, 384-well flat-bottom polystyrene plate (Thermo Fisher). 10 μL of serially diluted recombinant GABARAP or LC3B was added to each well. The plate was covered with foil and incubated with shaking for 1 h at room temperature. After incubation, the plate was read on a Tecan plate reader at $\lambda_{ex}$ = 494 nm and $\lambda_{em}$ = 519 nm. $K_d$ values were determined using curve fits using GraphPad Prism software as described,[2] using the curve fit equation below. Average $K_d$ and standard error of the mean were calculated using three independent trials.

$K_d$ curve fit equation: $y = m1 + (m2-m1) * (K_d+L+P)-((K_d+L+P)^2-4*L*P))^{0.5} / 2*P$

y is measured polarization, m1 is polarization at 0% bound, m2 is polarization at 100% bound, P is the total concentration of protein, and L is the total concentration of dye-labeled probe

#### AlphaScreen Assay

Recombinant GABARAP was diluted in assay buffer (25 mM HEPES, 150 mM NaCl, 0.1% Tween-20, 1 mg/mL BSA) to 50 nM and 5 μL was added in duplicate to a white, 384-well polystyrene plate (AlphaPlate, Revvity). Separate wells were prepared for serial dilutions of test peptides, acetylated-K1 (Ac-K1) as a positive control, and negative controls lacking K1 peptide or lacking GABARAP. A 25 μM Ac-K1 solution was prepared then serially diluted. 5 μL of serially diluted Ac-K1 was added to the Ac-K1 control row. 20 mM or 2 mM stocks of acetylated peptide in DMSO were used to prepare a 100 μM solution with either 2.5% or 5% DMSO, respectively, and then these solutions were serially diluted. 5 μL of diluted peptide solution was added to test wells. The plate was covered with foil, centrifuged at 1200 rpm for 3 min, and incubated at room temperature for 45 min. Biotinylated K1 (Bio-K1) was diluted to 50 nM and 5 μL was added to all wells except the appropriate negative control wells. The plate was covered with foil, centrifuged, and incubated for an additional 45 min. AlphaScreen streptavidin donor beads and nickel chelate acceptor beads (Revvity) were diluted to 100 μg/mL. 5 μL of acceptor beads were added to all wells. In the dark, 5 μL of donor beads were added to all wells. The plate was covered with foil, centrifuged, and incubated for an additional 1 h. The plate was read on a Tecan Spark plate reader (excitation 680 nM and emission 520-620 nM). Data were normalized using GraphPad Prism software using blank control wells as 0% bound and no inhibitor wells as 100% bound. After, the normalized data was fit with an $IC_{50}$ curve. Average $IC_{50}$ and standard error of the mean were calculated using three independent trials.

#### MDCK Permeability Assay

MDCK II (CRL-2936) cells were used in this permeability assay. Cells were cultured to ~70% confluency, trypsinized, and seeded on a Falcon 96-Multiwell Insert System (Corning 353938) at 2.5 x $10^5$ cells/mL. Once seeded, the cells were allowed to incubate for 5 days, washing the cells and replacing media on the second day of incubation. Cells were cultured in MEM with L-glutamine (Corning Inc., Corning, NY).

Prior to the assay, the growth medium was removed from both the donor chamber and Feeder Tray, and replaced with the transfer buffer, Hank's Balanced Salt Solution (HBSS) (Thermo

Fischer Scientific, Waltham, MA) to equilibrate the cells. During this time, the donor plate, a HTS 96 Square Well, Angled Bottom Plate (Corning 353925) was charged with 250 µL HBSS.

To initiate the assay, HBSS was removed from the donor chamber and replaced with 75 µL of test compound solution (10 µM, 5 µM Lucifer Yellow (LY) in HBSS). The Insert System was then placed into the Square Well Plate containing HBSS, covered with a lid, and incubated for 180 min (37 °C, 5% $CO_2$, 95% humidity). After the incubation period, the plates were separated and 100 µL of the acceptor solution was added to a fluorescence-compatible 96-well plate (Corning 3369). 150 µL of ACN was then added to the remaining 150 µL in the acceptor plate and 100 µL of this 1:1 solution was added to a 96 well assay plate (Corning 3357) for analysis by LC-MS. Mass identification and quantification used an inline Thermo Scientific Orbitrap VelosPro (FTMS mode). Samples were run in a minimum of triplicate with 10 µM carbamazepine as a control (Log($P_e$) = -4.14 cm/s)

The integrity of the cell monolayer was assessed by detection of LY in acceptor wells using a PerkinElmer Envision Reader (model 2105) equipped with the fluorescence intensity option (mirror: FITC D505 (PerkinElmer); excitation filter: Umbelliferone 460/25 (PerkinElmer); emission filter: FITC 535/25 (PerkinElmer).

Permeability rates across the cell monolayer were calculated via the following equation:

$$P_e = \frac{dM_r}{dt} * \frac{1}{Msa * C_D(0)}$$

Where:

$\frac{dMr}{dt}$ = flux of the compound across the cell monolayer

$Msa$ = surface area of filter membrane = 0.0804 $cm^2$

$C_D(0)$ = concentration of compound in the donor well at “t=0”

**Parallel Artificial Membrane Permeability Assay (PAMPA)**

Pure compound stocks (150 µL, 10 µM, 5% DMSO in PBS, pH 7.4) were added to a 96-well donor plate with 0.45 µm hydrophobic Immobilon-P membranes (Millipore MAIPNTR10) pre-soaked with a 1% solution of lecithin in n-dodecane. The donor plate was loaded into a 96-well Teflon acceptor plate (Millipore MSSACCEPTOR) with 300 µL of PBS with 5% DMSO. After 16-18 h, the acceptor and donor plates were separated, and 50 µL was collect from each well. Samples were diluted with 50 µL methanol and analyzed by LC-MS as described above. Samples were run in a minimum of triplicate with 10 µM carbamazepine as a control (Log($P_e$) = -4.97 cm/s). Exact time in seconds was used in the calculations.

$$P_e = -C * \ln\left(1 - \left(\frac{\%T}{100}\right)\right)$$

$$C = \frac{Vd * Va}{(Vd + Va) * Msa * Ts}$$

Permeation rates ($P_e$) were calculated from %T by the following equations:

$$\%T = \left(\frac{Ia}{[AnalyteEquil]}\right) * 100$$

$$AnalyteEquil = \frac{(Ia * Va) + (Id * Vd)}{Va + Vd}$$

Where:

Msa = Active surface area of membrane = 240 mm$^2$

Va = Volume of acceptor well= 300 µL

Vd = Volume of donor well= 150 µL

Ts = Assay run time (s)

Id = Donor intensity

Ia = Acceptor intensity

**X-ray crystallography**

The M15 peptide was dissolved in DMSO and mixed with GABARAP at a molar ratio of 3:2. After a 10-min incubation at RT, any occurring precipitate was removed by centrifugation (10 min at 20,000×*g* and 4 °C). The GABARAP–M15 complex was concentrated to a final protein concentration of 13–15 mg/mL using an Amicon Ultra-0.5 centrifugal filter with 3 kDa cutoff. Prior to application in crystallization trials, the protein-peptide complex was once again centrifuged for 30 min at 20,000×*g* and 4 °C. Search for crystallization conditions was performed by the sitting-drop vapor diffusion method using robotic systems Freedom Evo (Tecan) and Mosquito LCP (SPT Labtech) with commercially available screening sets. Experiments were set up by combining 200 nL of protein-peptide complex with 200 nL of reservoir solution, and plates were incubated at 20 °C.

Diffraction-quality samples used for X-ray structure determination occurred with a reservoir solution containing 0.2 M ammonium sulphate, 0.1 M sodium cacodylate trihydrate and 30% (w/v) PEG 8,000. The diffraction experiment was carried out at 100 K on beamline ID30A-3 of the European Synchrotron Radiation Facility (ESRF; Grenoble, France) tuned to a wavelength of 0.9677 Å, using an Eiger X 4M detector. Raw data (doi.org/10.15151/esrf-dc-2039871972) were processed using XDS and XSCALE,[4] including reflections up to a diffraction limit of 1.42 Å. Crystals belonged to space group I23. Initial phases were determined using the structure of GABARAP in complex with stapled peptide Pen8-*ortho* (PDB ID: 7ZL7) for molecular replacement in MOLREP,[5] revealing a single copy of the complex per asymmetric unit. A first refinement was done in REFMAC5 followed by evaluation in Coot.[6,7] In the course of alternating model building in Coot and reciprocal-space refinement in Phenix.refine,[8] the M15 peptide was added and the structure iteratively improved. Model evaluation was carried out with MolProbity[9] and the wwPDB validation system (https://validate-rcsb-2.wwpdb.org/), revealing good geometry without Ramachandran outliers or unusual rotamers. For full data collection and refinement statistics refer to Table S6. The final coordinates along with structure factor amplitudes have been uploaded to the wwPDB with ID 9I9X (doi.org/10.2210/pdb9I9X/pdb). In that PDB entry, peptide M15 is denoted as peptide IM-2.

### Molecular dynamics: Partial charges

Partial charges for norleucine and *N*-methylated norleucine were derived using the R.E.D. webserver.[10] The residues were capped by an acetyl group (ACE) at the N-terminus and an *N*-methyl group (NME) at the C-terminus. Two conformations were prepared for norleucine: one near the $\alpha_R$-helix with ($\phi$, $\psi$) values of (–60°, –40°) and one near the extended conformation with ($\phi$, $\psi$) values of (–130°, 120°). All amide bonds were in *trans* configuration. Four conformations were prepared for *N*-methylated norleucine, with ($\phi$, $\psi$) values of (–60°, –40°) or (–130°, 120°) and with the amide bond between ACE and *N*-methylated norleucine in *trans* configuration or *cis* configuration. The amide bond between the *N*-methylated norleucine and NME residue was kept in *trans* configuration. In these conformations, atomic partial charges were generated in accordance with the Amber99SB force field.[11] The "RESP-A1" charge model at the HF/6-31G* theory level was used to compute the charge set. The surface options in molecular electrostatic potential calculation were set to IOp(6/33=2,6/41=4,6/42=6). These conformations were optimized with frozen ($\phi$, $\psi$) values. During charge fitting, the charges of atoms belonging to the cap residues were restrained to the values in the Amber99SB force field. For norleucine, the partial charges for N and H were restrained to –0.4157 *e* and 0.2719 *e*, respectively. For both norleucine and *N*-methylated norleucine, the partial charges for C and O were restrained to 0.5973 *e* and –0.5679 *e*, respectively, consistent with the Amber99SB force field.[12] The scaling factor for 1,4 electrostatic energy was set to 0.8333, and the scaling factor for 1,4 van der Waals energy was set to 0.5000. The cap residues were removed from the molecule to create the designated molecular fragment. The derived partial charges for norleucine and *N*-methylated norleucine are shown in Fig. S1.

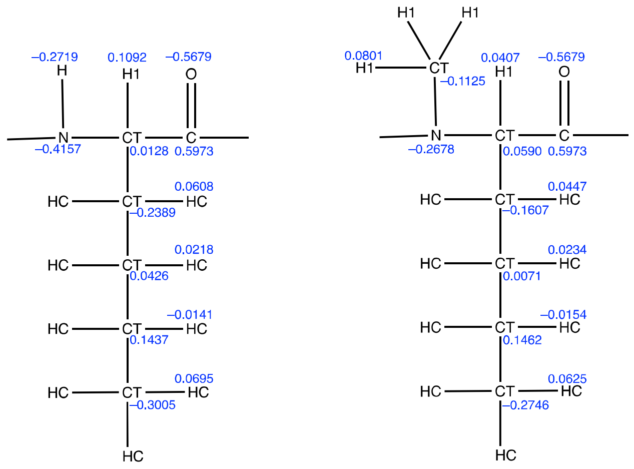

**Figure S1**. Atom types and partial charges for norleucine (left) and *N*-methylated norleucine (right).

### Molecular dynamics: Bias-exchange metadynamics (BE-META) simulations

The peptides were subjected to simulation using the BE-META method[13] within the RSFF2 force field[14] along with the TIP3P water model. The parameters for the linker were adopted from a previous paper.[15] The BE-META simulations were conducted using GROMACS 2018.6[16] patched with the PLUMED 2.5.1 plugin.[17]

Two independent simulations, initiated with distinct initial structures (S1 and S2) for each peptide, were executed to assess the convergence of simulation outcomes. For the linear, unstapled peptides, eight biased replicas were employed, biasing ($\phi_1$,$\psi_1$), ($\phi_2$,$\psi_2$), ($\phi_3$,$\psi_3$), ($\phi_4$,$\psi_4$), $\psi_5$, ($\psi_1$, $\phi_2$), ($\psi_2$, $\phi_3$), and ($\psi_3$, $\phi_4$), respectively. For the stapled peptides, seventeen biased replicas were employed, biasing ($d_2$, $d_1$), ($d_1$, $\chi_{2,2}$), ($\chi_{2,2}$, $\chi_{2,1}$), ($\chi_{2,1}$, $\psi_2$'), ($\psi_2$, $\phi_3$), ($\phi_3$, $\psi_3$), ($\psi_3$, $\phi_4$), ($\phi_4$', $\chi_{4,1}$), ($\chi_{4,1}$, $\chi_{4,2}$), ($\chi_{4,2}$, $d_4$), ($d_4$, $d_3$), ($d_3$, $d_2$), ($\chi_{2,1}$, $\phi_2$), ($\psi_4$, $\chi_{4,1}$), ($\phi_1$, $\psi_1$), ($\psi_1$, $\phi_2$), and $\psi_5$, respectively (see Fig. S2 for dihedral definitions). Additionally, five neutral replicas (i.e., replicas with no bias) were incorporated to obtain an unbiased structural ensemble for subsequent analysis. Exchange attempts were made between different replicas every 5 ps. The simulation lengths were 200 ns.

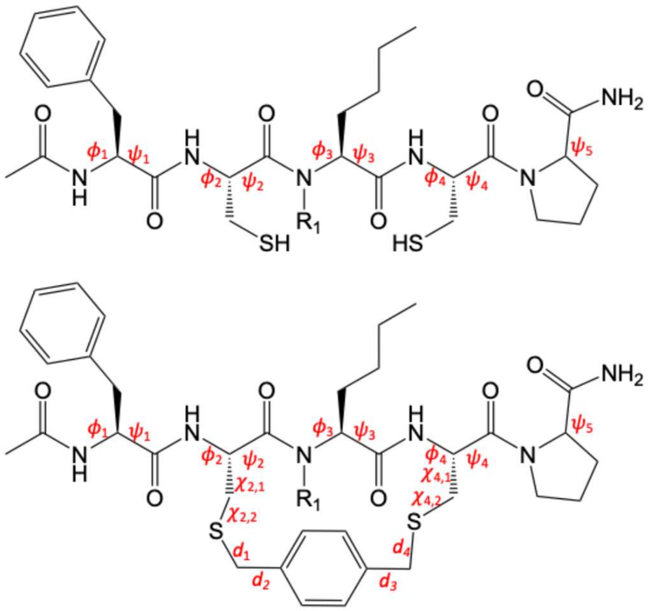

**Figure S2**. Definition of the dihedrals used in the BE-META simulations. $R_1$ denotes either an amide proton for non-*N*-methylated amino acids or a methyl group for *N*-methylated amino acids.

### Principal component analysis

To evaluate the convergence of the two distinct sets of simulations originating from initial structures S1 and S2, dihedral principal component analysis (dPCA) was conducted on the backbone ($\phi$, $\psi$) dihedrals.[18,19] The last 100 ns of the trajectories from the neutral replicas were used for analysis. dPCA was performed on frames from both S1 and S2 simulations. After obtaining the common principal components (PCs), frames from S1 and S2 simulations were individually projected into the 3D space defined by the top three PCs. Subsequently, a 3D density distribution was calculated for each simulation (S1 and S2). To assess convergence, specifically the similarity between the S1 and S2 density distributions, the normalized integrated product (NIP) was computed.[20] The NIP value ranges from 0 (indicating no overlap) to 1.0 (indicating perfect similarity). The NIP values between S1 and S2 for all peptides were above 0.9, indicating the simulations had converged. Upon confirming convergence, a final analysis was performed using trajectories from the last 100 ns of the neutral replicas.

### Supplemental Data

**Table S1. Expected and observed masses of fluorescein-tagged peptides used in fluorescence polarization assays.** All peptides were amidated at the C-terminus. The N-terminus was fluorescein-tagged with a linker of two β-alanine residues. Underlined residues are *N*-methylated. J denotes norleucine and X denotes penicillamine. dmb refers to dimethylbenzene.

| Name | Sequence | Modification | Expected Mass | Observed Mass |
|---|---|---|---|---|
| M1 | TDDFVJVPA | none | 1475.57 | 1497.97 [+Na] |
| M2 | TD**C**FV**C**VPA | allyl | 1533.73 | 1555.73 [+Na] |
| M3 | TD**C**FV**C**VPA | *para*-dmb | 1555.74 | 1554.81, 1576.89 [+Na] |
| M4 | TD**C**F**C**JVPA | allyl | 1547.76 | 1545.92, 1567.76 [+Na] |
| M5 | TD**C**F**C**JVPA | *para*-dmb | 1569.77 | 1567.94 |
| M6 | TDDF**C**J**C**PA | allyl | 1563.72 | 1561.47, 1583.37 [+Na] |
| M7 | TDDF**C**J**C**PA | *meta*-dmb | 1585.72 | 1583.11, 1605 [+Na] |
| M8 | TDDF**C**J**C**PA | *para*-dmb | 1585.72 | 1584.22 |
| M9 | TDDFVJVPA | *N*-methyl at Phe4 | 1489.60 | 1487.73, 1510.50 [+Na] |
| M10 | TDDFVJVPA | *N*-methyl at Val5 | 1489.60 | 1489.37, 1512.08 [+Na] |
| M11 | TDDFVJVPA | *N*-methyl at Nle6 | 1489.60 | 1488.41, 1510.22 [+Na] |
| M12 | TDDFVJVPA | *N*-methyl at Val7 | 1489.60 | 1508.88 [+Na] |
| Mpen1 | TDDF**X**J**C**PA | allyl | 1591.77 | 1592.93 [+H] |
| Mpen2 | TDDF**X**J**C**PA | *para*-dmb | 1613.78 | 1613.84, 1636.28 [+Na] |
| Mpen3 | TDDF**C**J**X**PA | allyl | 1591.77 | 1591.52, 1613.38 [+Na] |
| Mpen4 | TDDF**C**J**X**PA | *para*-dmb | 1613.78 | 1614.98 [+H], 1637.09 [+Na] |
| Mpen5 | TDDF**X**J**X**PA | allyl | 1619.82 | 1618.45, 1641.32 [+Na] |
| Mpen6 | TDDF**X**J**X**PA | *para*-dmb | 1641.83 | 1642.55, 1664.53 [+Na] |
| M13 | TDD**W**VJVPA | Phe4 to Trp (retained below) | 1514.61 | 1515.10 [+H], 1537 [+Na] |
| M14 | TDD**WC**J**C**PA | *para*-dmb | 1624.76 | 1621.02, 1643.70 [+Na] |
| M15 | TDD**W**VJVPA | *N*-methyl Nle6 | 1528.64 | 1528.52 |
| M16 | TDD**WC**J**C**PA | *N*-methyl Nle6, allyl | 1616.78 | 1619.49 [+3H], 1642.14 [+Na] |
| M17 | TDD**WC**J**C**PA | *N*-methyl Nle6, *para*-dmb | 1638.79 | 1638.84, 1660.74 [+Na] |
| M18 | **WC**J**C**PA | *para*-dmb, N-term truncation | 1293.48 | 1294.26 [+H], 1317.30 [+Na] |
| M19 | TDD**WC**J**C** | *para*-dmb, C-term truncation | 1456.56 | 1454.59, 1476.55 [+Na] |
| M20 | **WC**J**C** | *para*-dmb, full truncation | 1125.28 | 1125.38 |
| M21 | **W**VJVPA | *N*-methyl Nle6, N-term truncation | 1197.36 | 1198.22, 1221.14 [+Na] |
| M22 | TDD**W**VJV | *N*-methyl Nle6, C-term truncation | 1360.44 | 1359.03, 1381.87 [+Na] |
| M23 | **W**VJV | *N*-methyl Nle6, full truncation | 1029.16 | 1030.46, 1052.44 [+Na] |

**Table S2. Expected and observed masses of acetylated peptides used in AlphaScreen assays.** All peptides were amidated at the C-terminus. All peptides were acetylated at the N-terminus, except for M24 which was capped with 3-indole-acetic acid (denoted with Θ). Underlined residues are *N*-methylated. J denotes norleucine and dmb refers to dimethylbenzene.

| Name | Sequence | Modification | Expected Mass | Observed Mass |
|---|---|---|---|---|
| M13 | TDD**W**VJVPA | None | 1056.19 | 1077.10 [+Na] |
| M14 | TDD**WC**J**C**PA | *para*-dmb | 1166.33 | 1168.42 [+H] |
| M15 | TDD**W**VJVPA | *N*-methyl Nle6 | 1070.21 | 1093.09 [+Na] |
| M23 | **W**VJV | *N*-methyl Nle6 | 570.74 | 593.76 [+Na] |
| M24 | **Θ**VJV | *N*-methyl Nle6,<br>3-indole acetic acid cap | 513.68 | 514.18 [+H] |

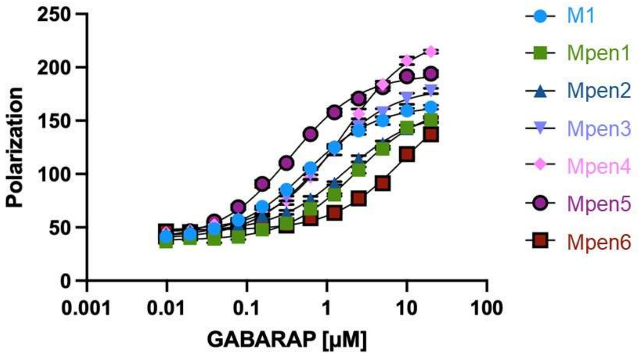


| Peptide | Sequence | Modification | $K_d$ (nM) |
|---|---|---|---|
| M1 | TDDFVJVPA | none | 557 ± 22 |
| Mpen1 | TDDF**X**J**C**PA | allyl | 2310 ± 50 |
| Mpen2 | TDDF**X**J**C**PA | *para*-dmb | 1720 ± 190 |
| Mpen3 | TDDF**C**J**X**PA | allyl | 865 ± 72 |
| Mpen4 | TDDF**C**J**X**PA | *para*-dmb | 1640 ± 110 |
| Mpen5 | TDDF**X**J**X**PA | allyl | 350 ± 4 |
| Mpen6 | TDDF**X**J**X**PA | *para*-dmb | > 8071 |

**Figure S3.** Fluorescence polarization data and curve fits for penicillamine-substituted stapled peptides binding to recombinant GABARAP. **X** denotes penicillamine and **J** denotes norleucine.

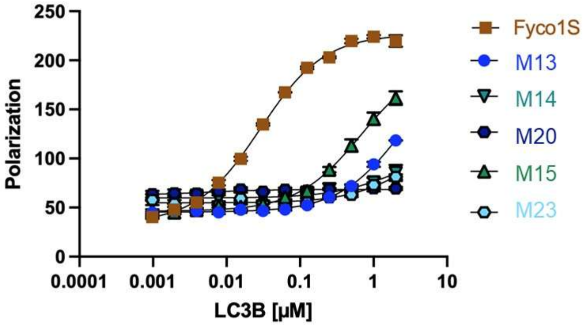


| Peptide | Sequence | Modification | $K_d$ (nM) |
|---|---|---|---|
| Fyco1S | DDAVFDIITDEELCQIQEW | (positive control) | 31.5 ± 2.0 |
| M13 | TDD**W**VJVPA | F4W | > 2534 |
| M14 | TDD**WC**J**C**PA | F4W, *para*-dmb | No binding |
| M20 | **WC**J**C** | F4W, *para*-dmb, full truncation | No binding |
| M15 | TDD**W**V<u>J</u>VPA | F4W, *N*-Me Nle6 | > 2211 |
| M23 | **W**V<u>J</u>V | F4W, *N*-Me Nle6, full truncation | No binding |

**Figure S4.** Fluorescence polarization data and curve fits for selected peptides binding to recombinant LC3B. Underlined residues are *N*-methylated. **J** denotes norleucine. Fyco1S is an LC3B-binding control peptide reported previously.[1,2]

**Table S3. Equilibrium dissociation constants ($K_d$, nM) for fluorescein-tagged peptides in the fluorescence polarization assay with recombinant GABARAP.** $K_d$ values correspond to curve fits shown in Figure S5. For curve fits where saturation at high concentrations of GABARAP was not observed, $K_d$ values from curve fits are interpreted as the minimum likely $K_d$ value indicated by the data.

| Peptide | Trial 1 | Trial 2 | Trial 3 | Average | SEM |
|---|---|---|---|---|---|
| M1 | 557 | 519 | 594 | 557 | 22 |
| M2 | 2251 | 2359 | 2615 | 2408 | 110 |
| M3 | >26441 | >17298 | >27640 | >23793 | |
| M4 | >6090 | >5617 | >5681 | >5796 | |
| M5 | >24563 | >21295 | >11876 | >19245 | |
| M6 | 2630 | 2020 | 2162 | 2271 | 180 |
| M7 | 2935 | 2746 | 2867 | 2849 | 60 |
| M8 | 236 | 178 | 239 | 218 | 20 |
| M9 | >6277 | >7415 | >5882 | >6525 | |
| M10 | No binding | No binding | No binding | No binding | |
| M11 | 104 | 100 | 118 | 107 | 3.06 |
| M12 | >10567 | >21450 | >30762 | >20926 | |
| Mpen1 | 2411 | 2232 | 2275 | 2306 | 54 |
| Mpen2 | 1655 | 1428 | 2074 | 1719 | 189 |
| Mpen3 | 733 | 981 | 880 | 865 | 72 |
| Mpen4 | 1443 | 1668 | 1800 | 1637 | 110 |
| Mpen5 | 336 | 340 | 373 | 350 | 4 |
| Mpen6 | >9849 | >8231 | >8024 | >8071 | |
| M13 | 36.8 | 47.5 | 45.6 | 43.3 | 3 |
| M14 | 13.7 | 12.1 | 18.4 | 14.7 | 2 |
| M15 | 9.08 | 10.4 | 9.45 | 9.64 | 0.4 |
| M16 | 175 | 165 | 141 | 160 | 10 |
| M17 | 81.0 | 48.3 | 84.4 | 71.2 | 12 |
| M18 | 254 | 191 | 200 | 215 | 20 |
| M19 | 283 | 282 | 256 | 274 | 9 |
| M20 | >1719 | >2144 | >1116 | >1660 | |
| M21 | 119 | 142 | 125 | 129 | 7 |
| M22 | 22.7 | 19.9 | 25.0 | 22.5 | 2 |
| M23 | 175 | 155 | 157 | 162 | 6 |

**Figure S5. Complete data sets and curve fits for fluorescein-tagged peptides in the fluorescence polarization assay with recombinant GABARAP.** Three independent trials are shown. Error bars, when visible, show standard error of the mean of three technical replicates within each trial.

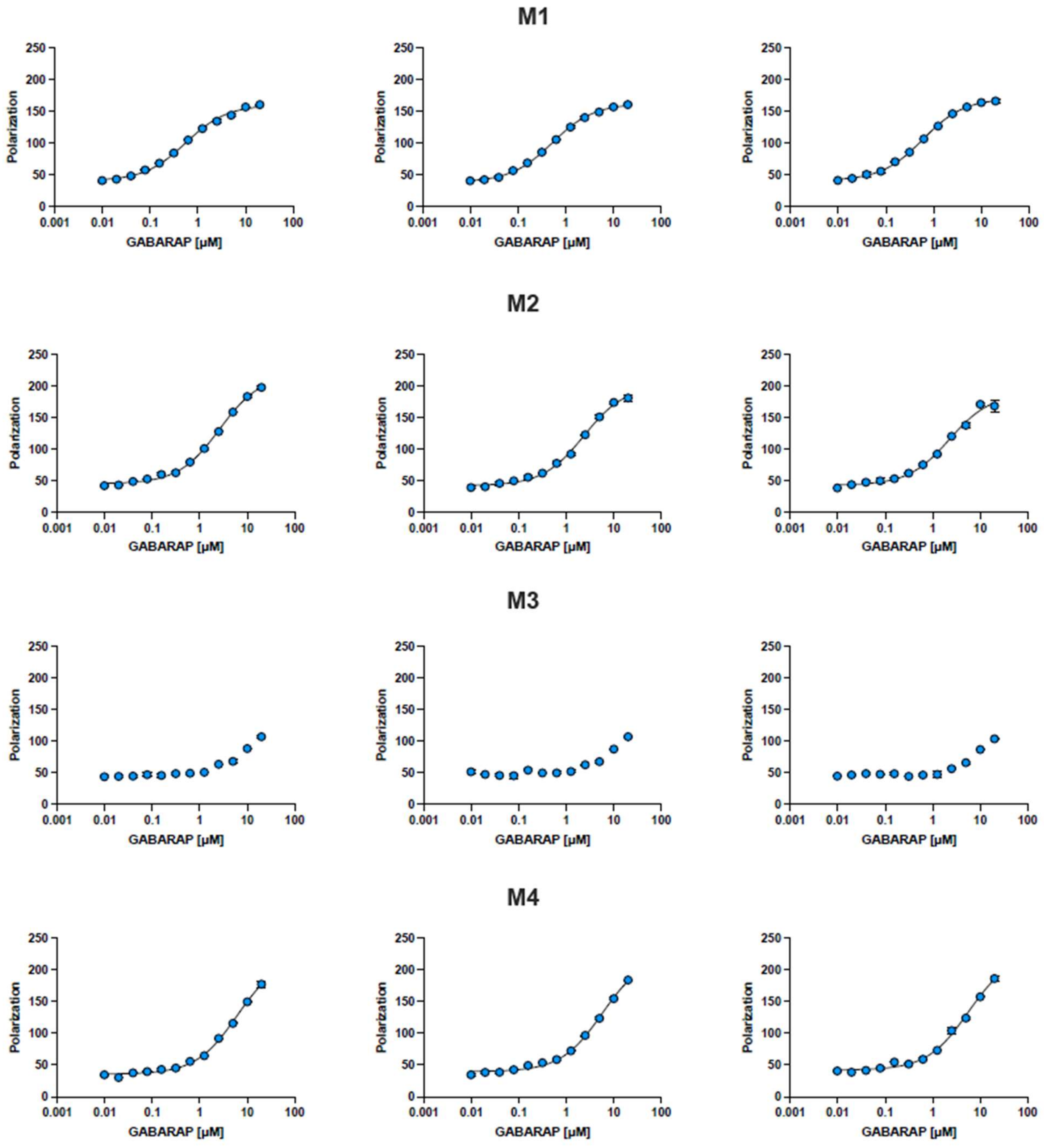

## M5

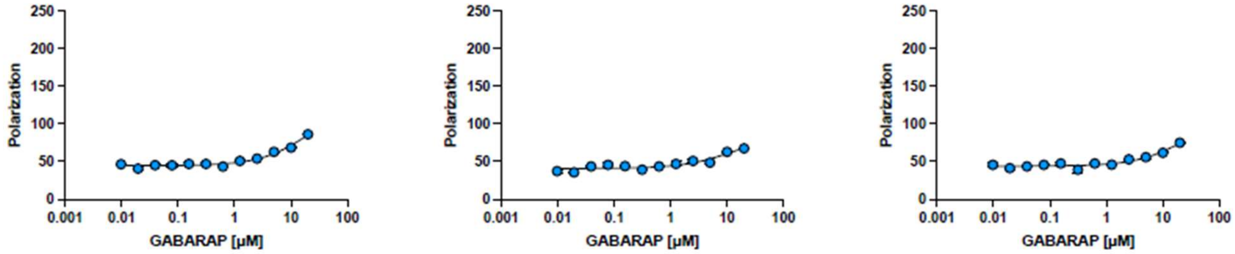
Polarization
GABARAP [µM]

## M6

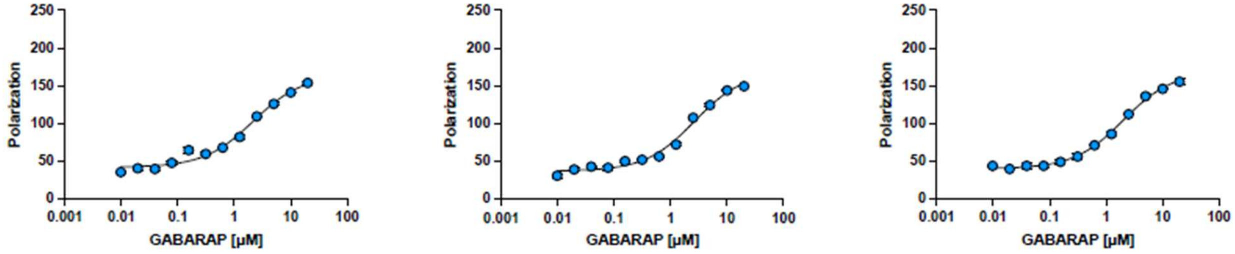
Polarization
GABARAP [µM]

## M7

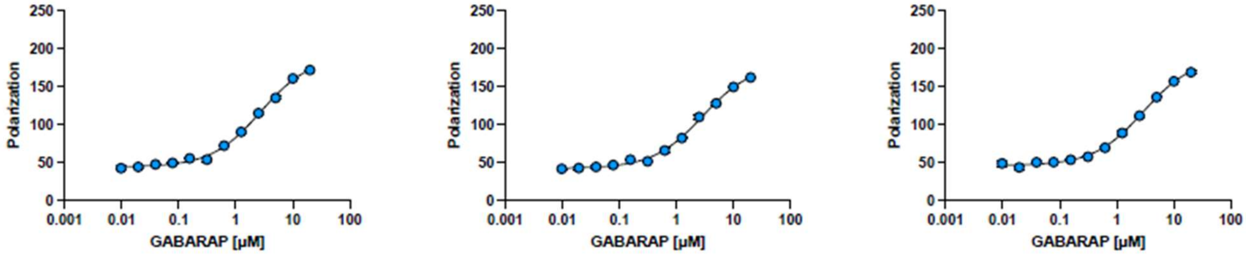
Polarization
GABARAP [µM]

## M8

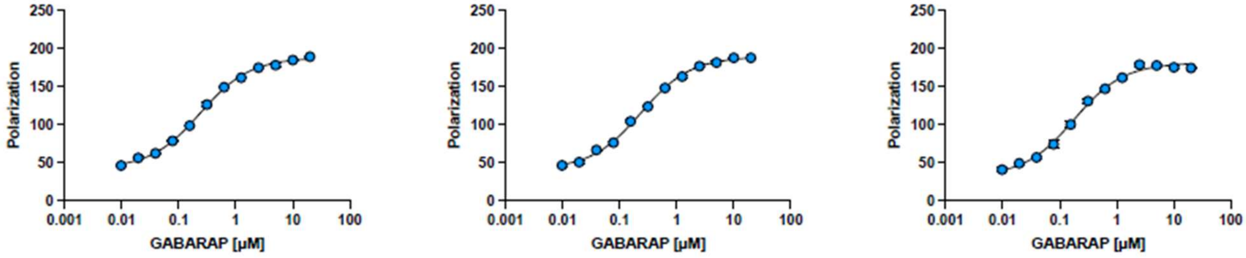
Polarization
GABARAP [µM]

## M9

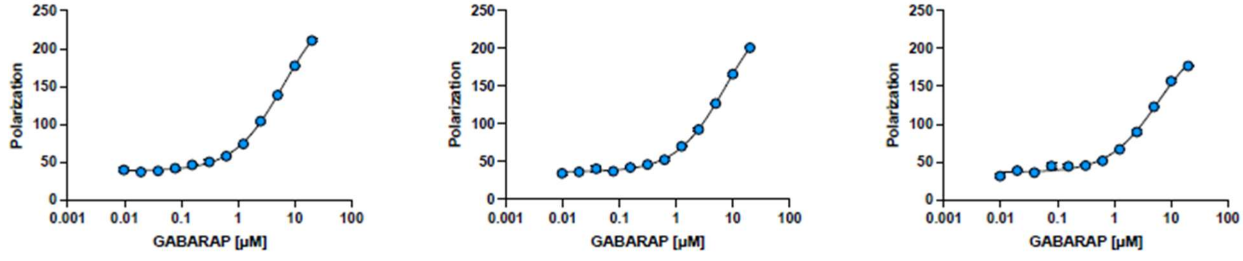
Polarization
GABARAP [µM]

**M10**

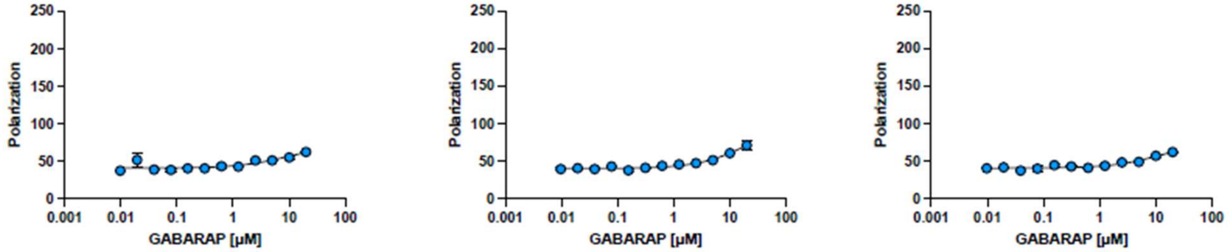


**M11**

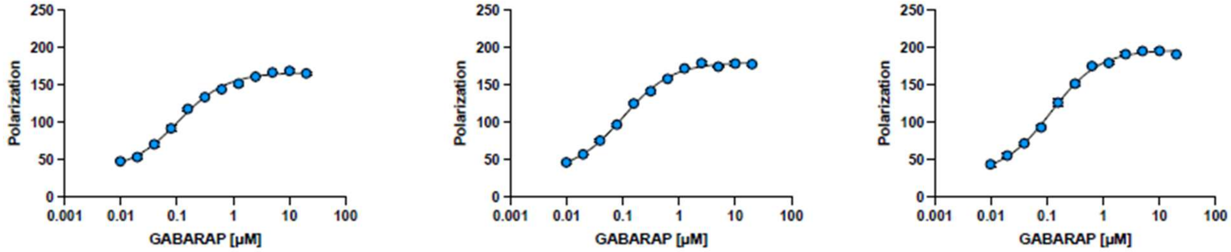


**M12**

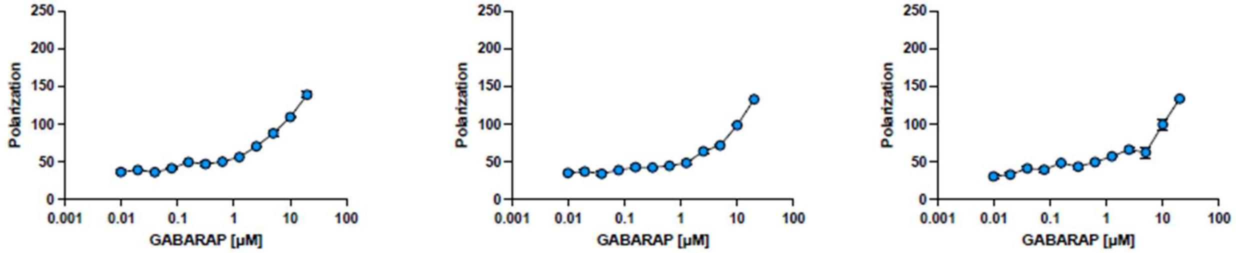


**Mpen1**

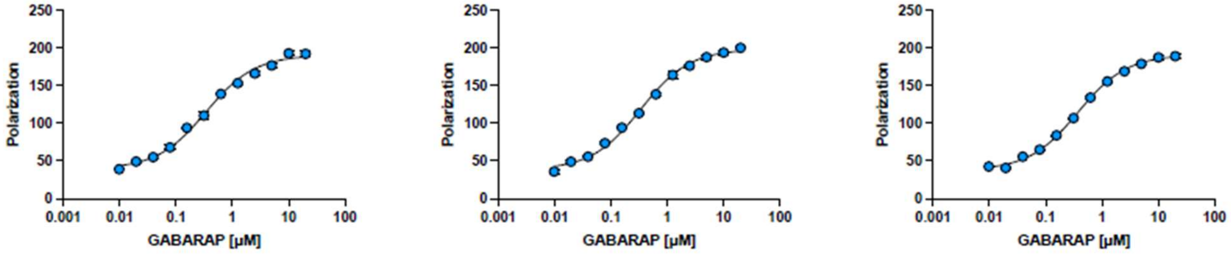


**Mpen2**

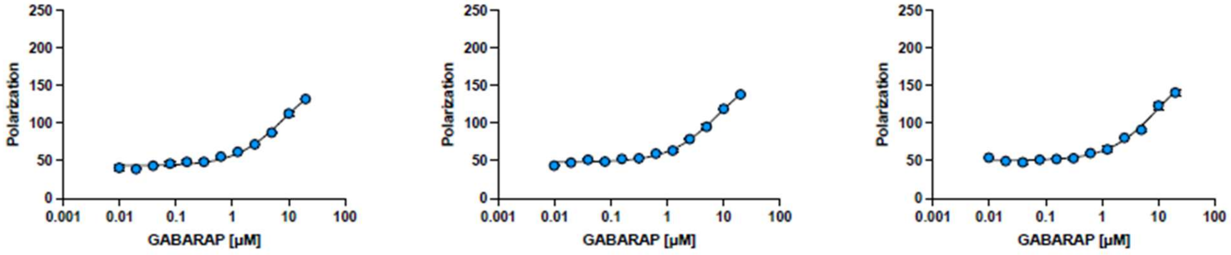

### Mpen3

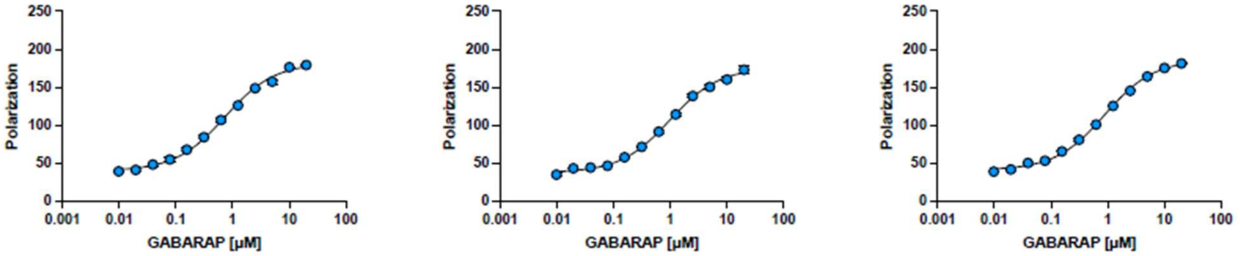


### Mpen4

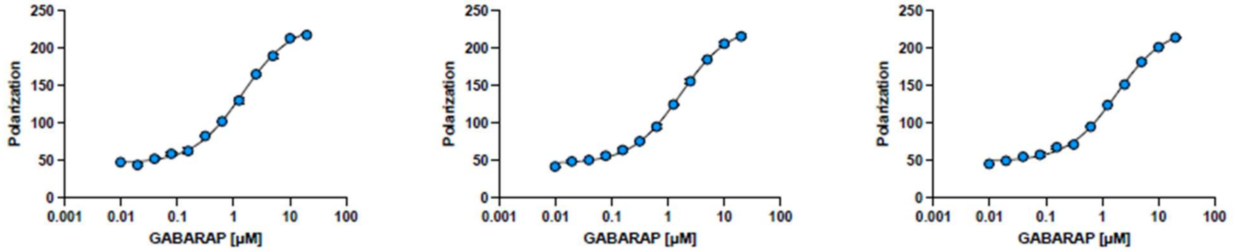


### Mpen5

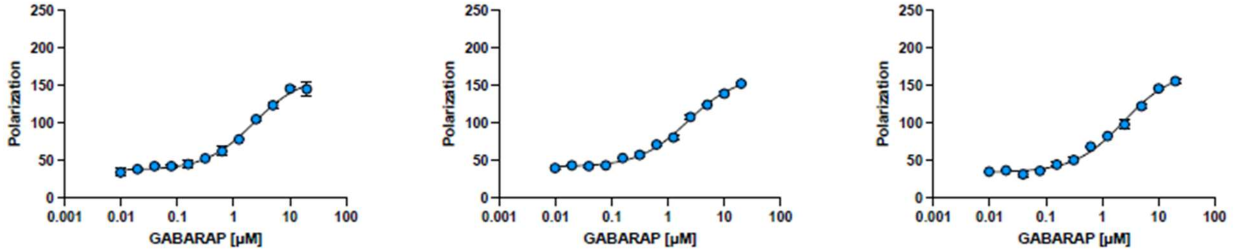


### Mpen6

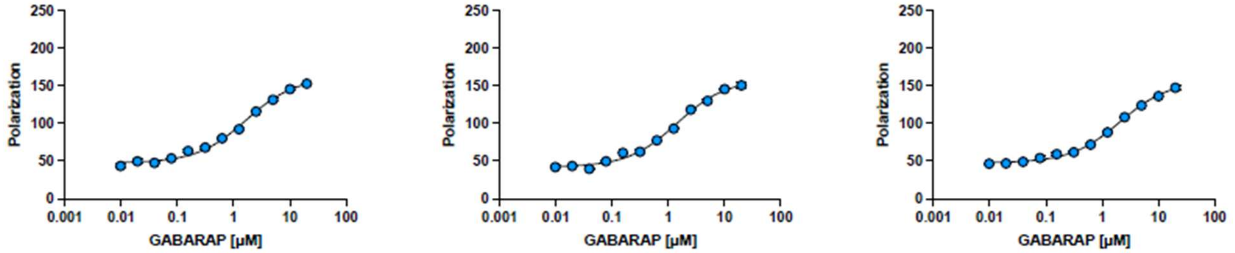


## M13

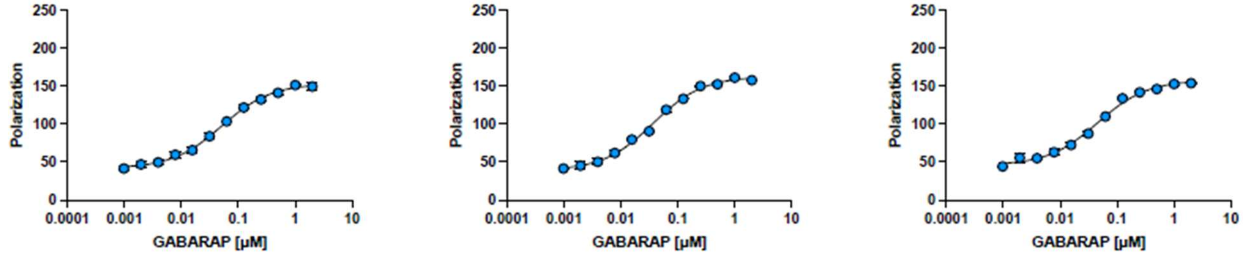

## M14

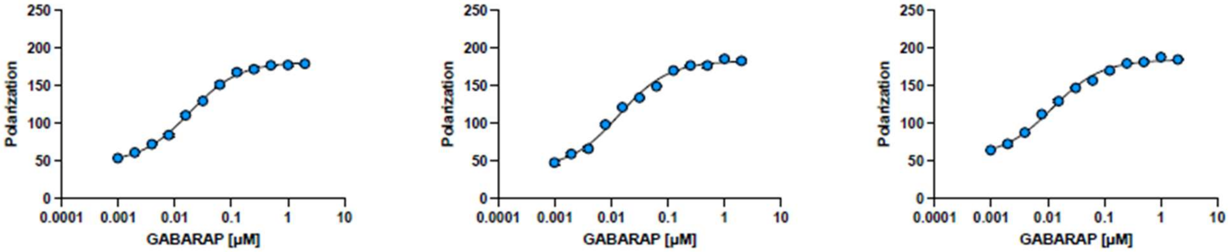


## M15

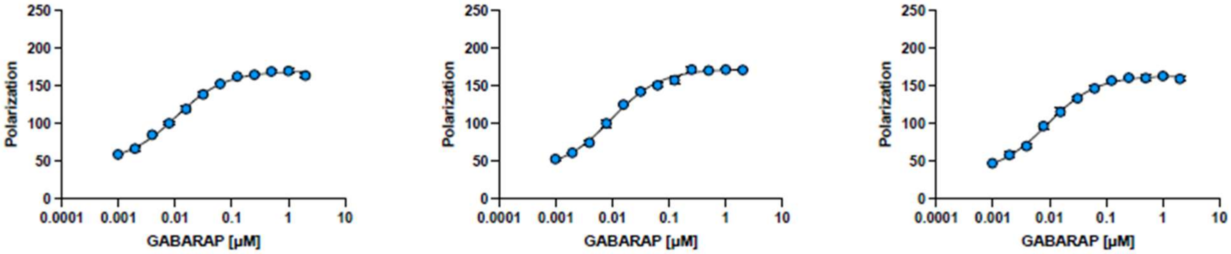


## M16

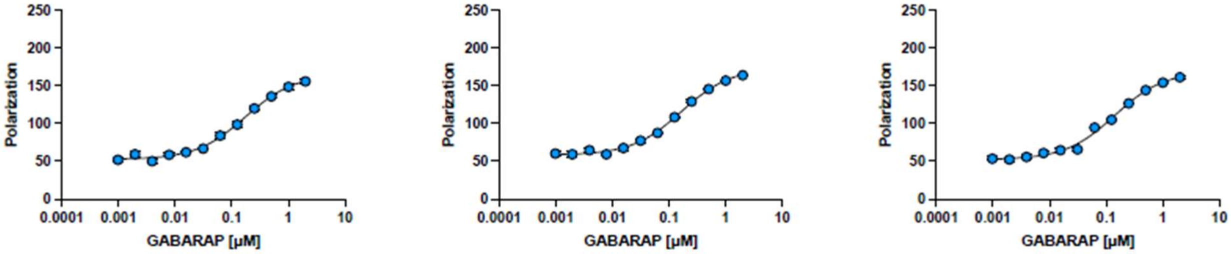


## M17

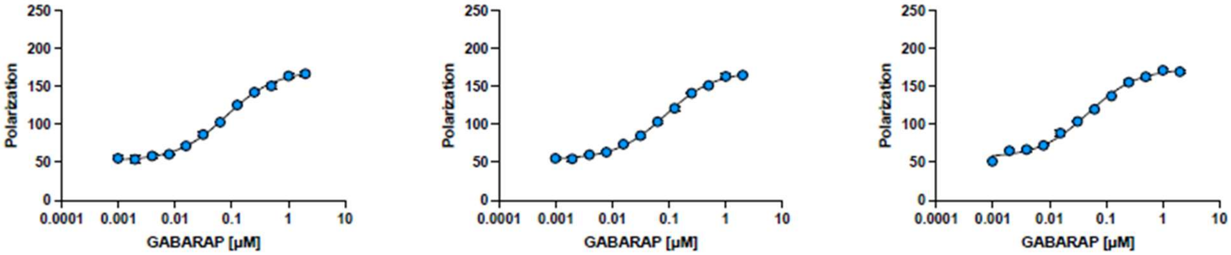


## M18

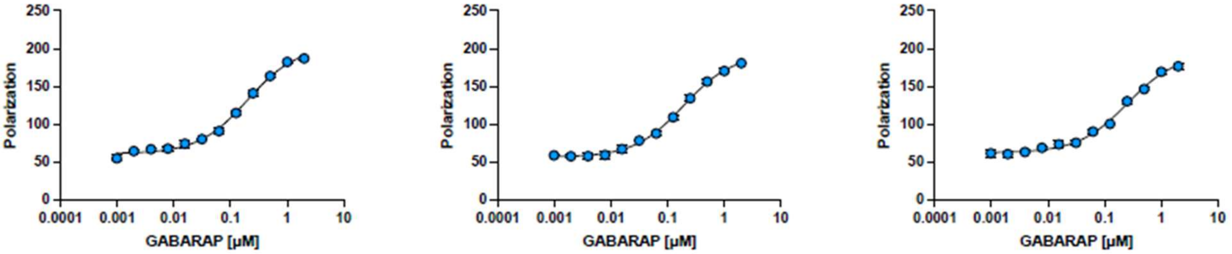

## M19

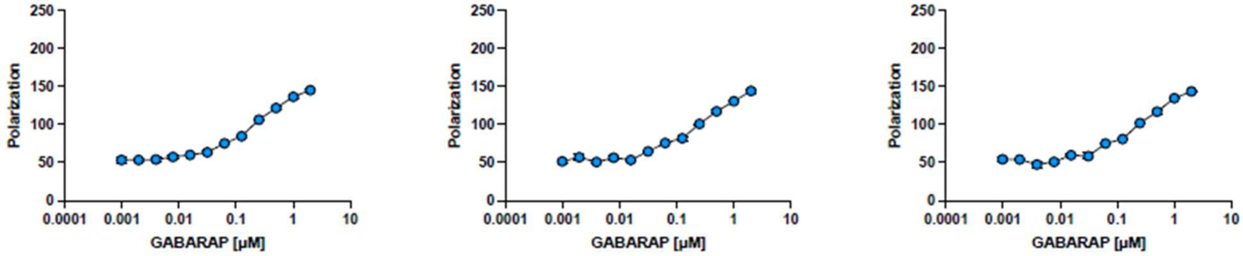


## M20

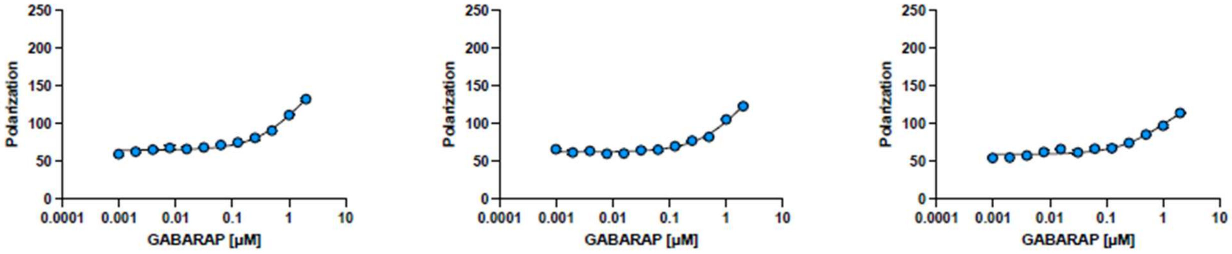


## M21

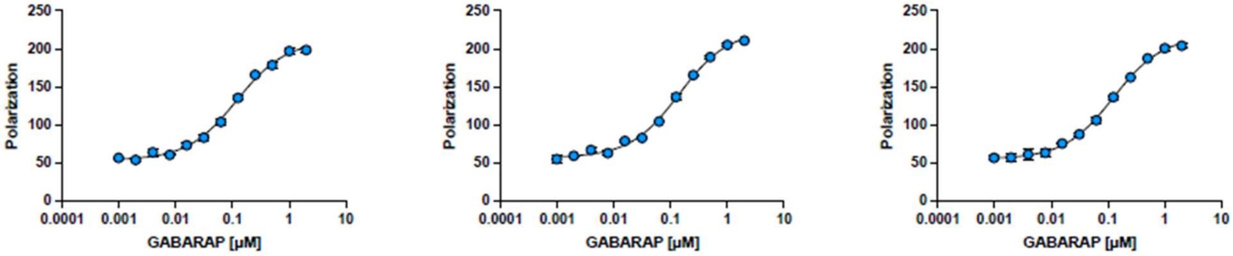


## M22

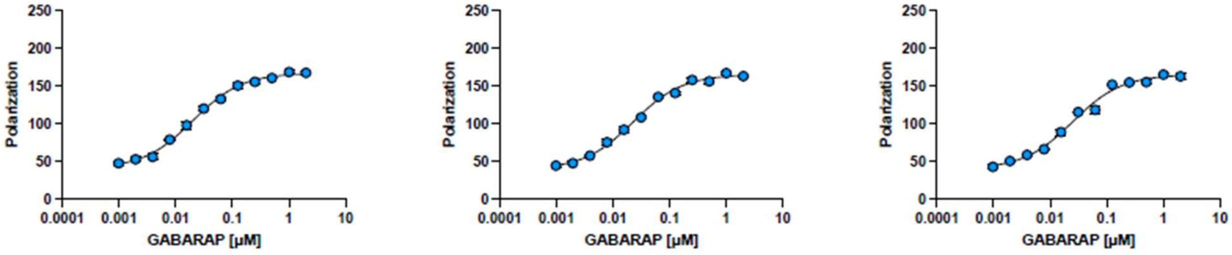


## M23

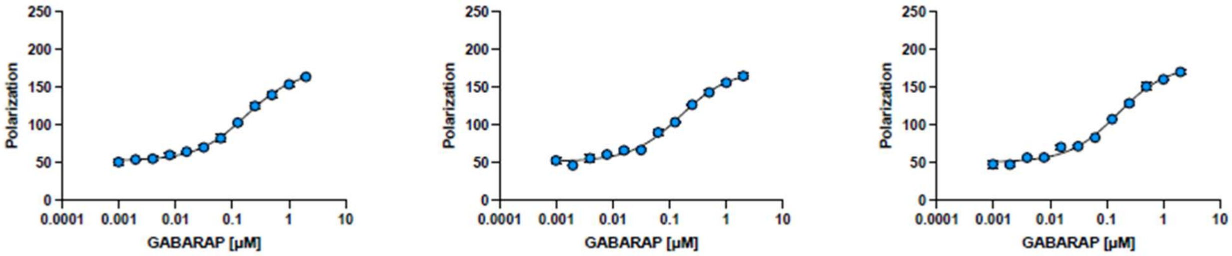

### Simulation results

Ramachandran plots showing the ensembles of each peptide are shown in main text Fig. 3. Comparing the unmodified peptide to the stapled peptide, we observed that the staple generally moves the backbone dihedrals towards the upper-left "extended" β-strand region of the Ramachandran plot; this effect was particularly noticeable for the norleucine residue (main text Fig. 3). Comparing the unmodified peptide to the *N*-methylated peptide, we observed a shift towards positive $\phi$ values at the norleucine residue. This result is somewhat counterintuitive, as *N*-methylated L-amino acids are generally expected to still favor negative $\phi$ values. To test whether this observation was due to a force field artifact, we examined the conformational ensemble of *N*-methylated alanine dipeptide using several force fields: RSFF2,[14] Amber14SB,[21] Amber19SB,[22] OPLS2005,[23] and OpenFF.[24] In all simulations, the $\phi$ angles predominantly favored positive values. Finally, for the peptide which was both stapled and *N*-methylated, the $\phi$ values at the central residue shifted back towards negative values, with the dihedral angle distributions lying between the PPII and α-helix regions (main text Fig. 3).

**Table S4. Inhibitory constants ($IC_{50}$, nM) of acetylated peptides in the AlphaScreen assay with recombinant GABARAP and biotinylated peptide K1.** $K_d$ values correspond to curve fits shown in Figure S6.

| Peptide | Trial 1 | Trial 2 | Trial 3 | Average | SEM |
|---|---|---|---|---|---|
| M13 | 688 | 530 | 339 | 519 | 101 |
| M14 | 378 | 457 | 417 | 418 | 23 |
| M15 | 318 | 230 | 221 | 256 | 31 |
| M23 | 5375 | 1806 | 3816 | 3666 | 1003 |
| M24 | 18973 | 16828 | 14340 | 16714 | 1339 |

**Figure S6. Complete data sets and curve fits for acetylated peptides in the AlphaScreen assay with recombinant GABARAP and biotinylated peptide K1.** Three independent trials are shown. Error bars, when visible, show standard error of the mean of three technical replicates within each trial.

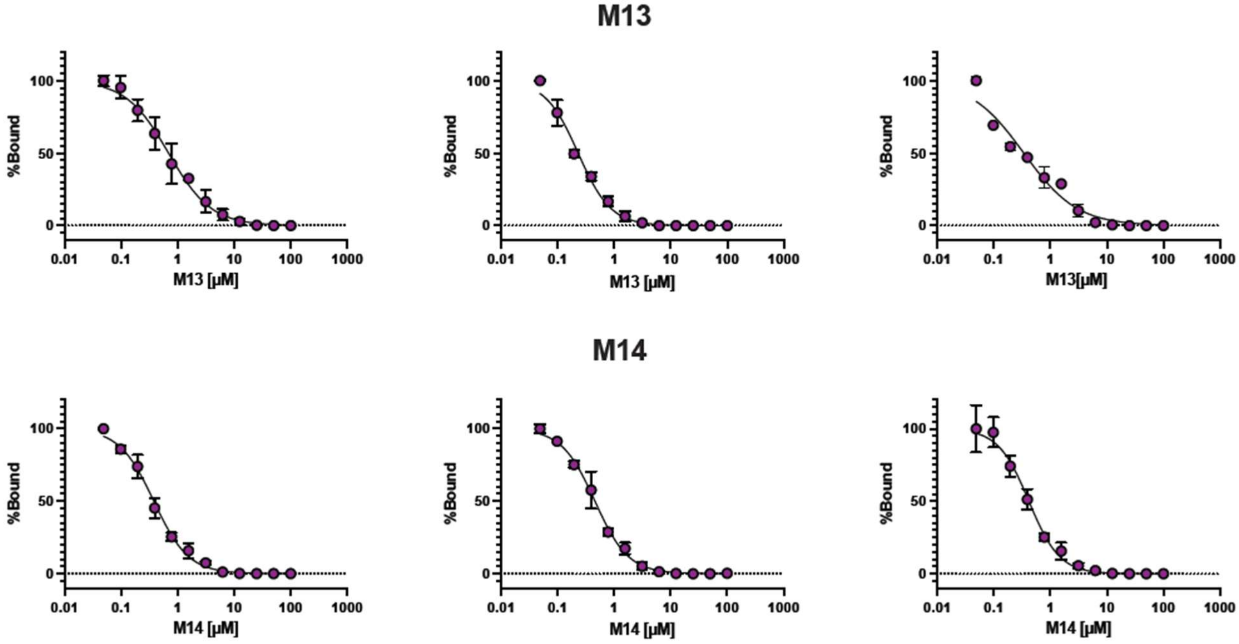

**M15**

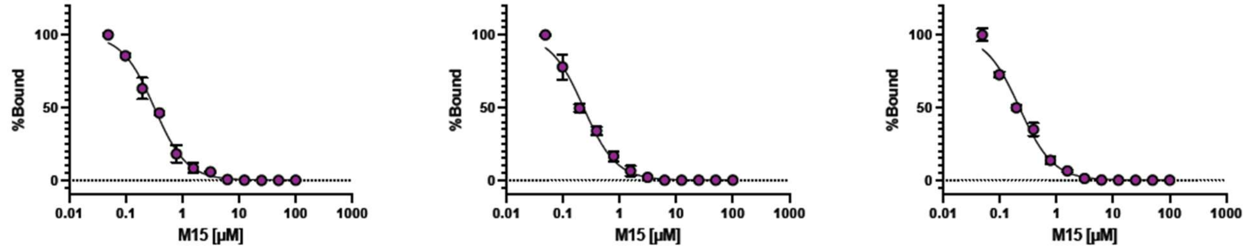


**M23**

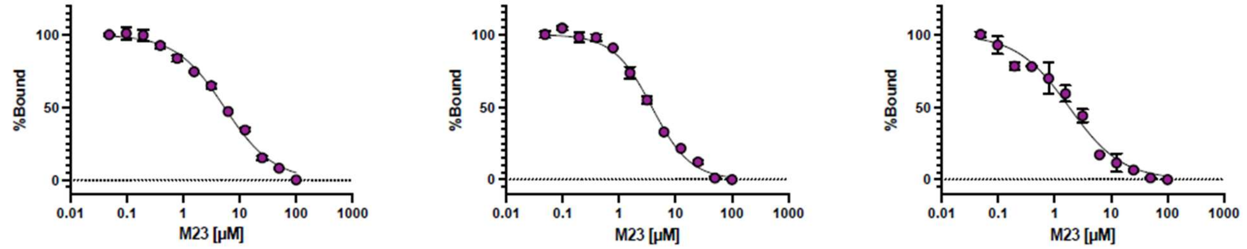


**M24**

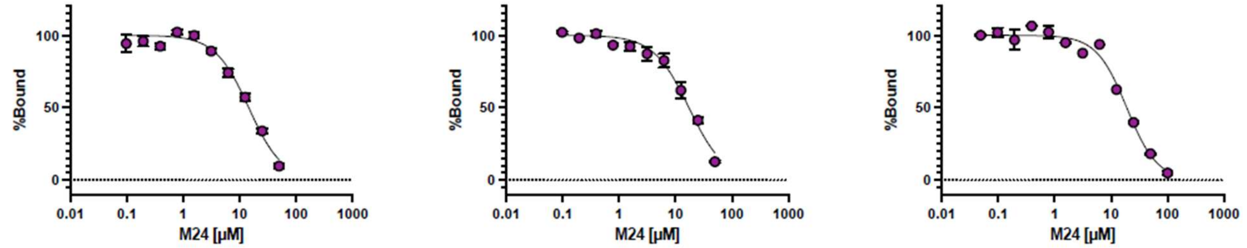


**Table S5. Equilibrium dissociation constants ($K_d$, nM) for fluorescein-tagged peptides in the fluorescence polarization assay with recombinant LC3B.** $K_d$ values correspond to curve fits shown in Figure S7. For curve fits where saturation at high concentrations of LC3B was not observed, $K_d$ values from curve fits are interpreted as the minimum likely $K_d$ value indicated by the data.

| Peptide | Trial 1 | Trial 2 | Trial 3 | Average | SEM |
|---|---|---|---|---|---|
| FYCO1S | 34.7 | 28.8 | 30.9 | 31.5 | 2 |
| M13 | > 2369 | > 2433 | > 2800 | > 2534 | n/a |
| M14 | No binding | No binding | No binding | No binding | n/a |
| M20 | No binding | No binding | No binding | No binding | n/a |
| M15 | > 1324 | > 3508 | > 1800 | > 2211 | n/a |
| M23 | No binding | No binding | No binding | No binding | n/a |

**Figure S7. Complete data sets and curve fits for fluorescein-tagged peptides in the fluorescence polarization assay with recombinant LC3B.** Three independent trials are shown. Error bars, when visible, show standard error of the mean of three technical replicates within each trial.

**FYCO1S**

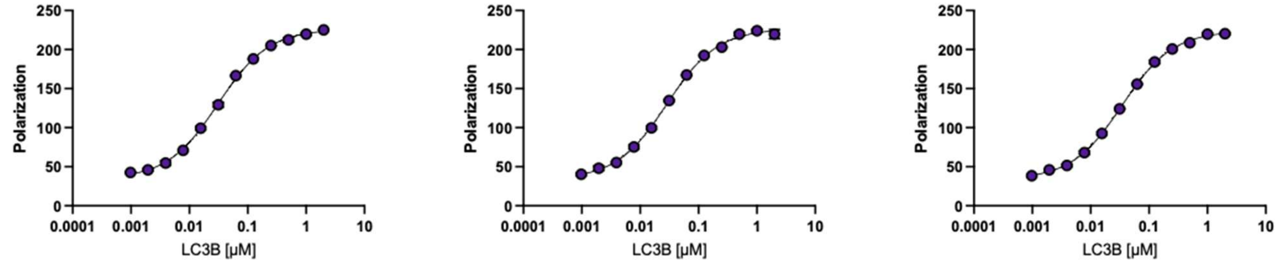


**M13**

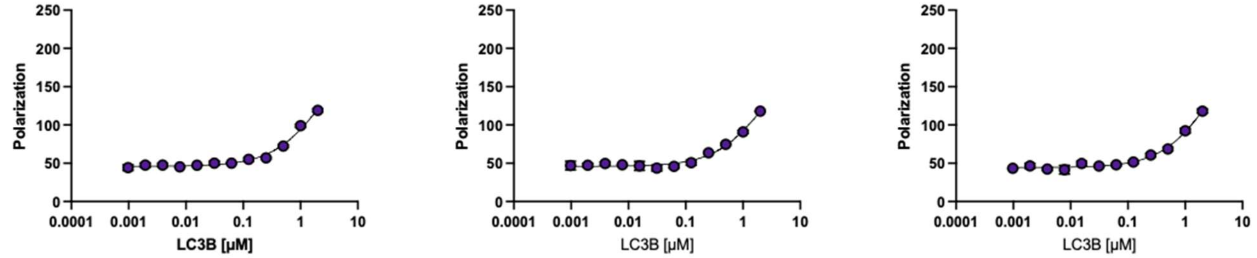


**M14**

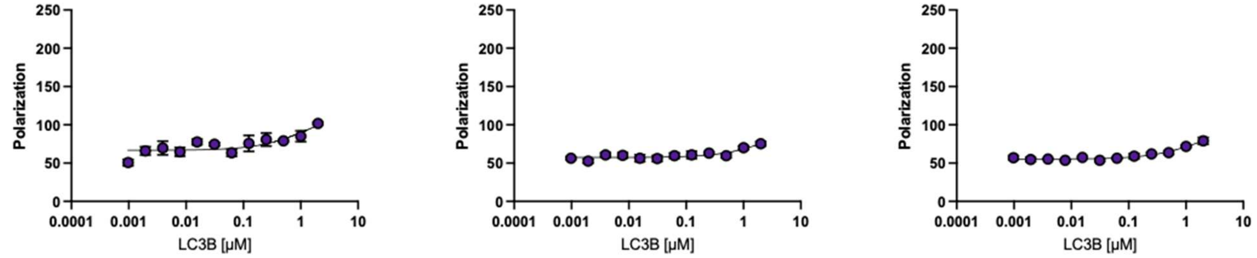


**M15**

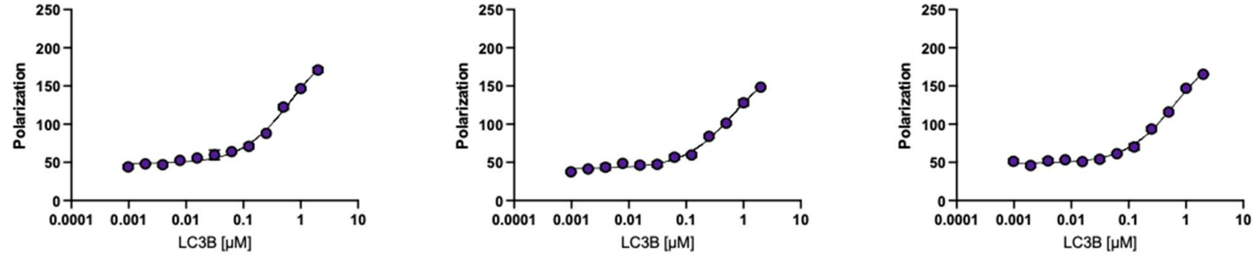

**M20**

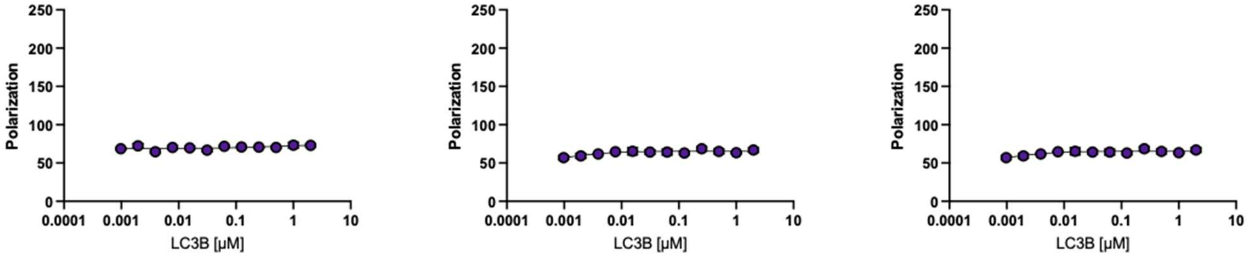
Polarization
0
50
100
150
200
250
0.0001
0.001
0.01
0.1
1
10
LC3B [μM]

**M23**

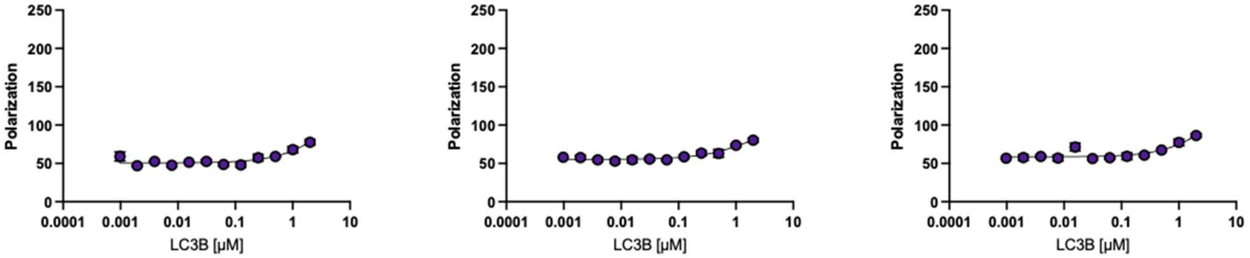
Polarization
0
50
100
150
200
250
0.0001
0.001
0.01
0.1
1
10
LC3B [μM]

**Table S6. Data collection and refinement statistics for the GABARAP–M15 crystal structure.** Values in parentheses refer to the outer (highest-resolution) shell.

| | **GABARAP–M15**<br>**PDB ID 9I9X** |
|---|---|
| **Data collection** | |
| **Resolution range (Å)** | 41.37–1.42 (1.46–1.42) |
| **Space group** | I 2 3 |
| **Unit cell: a, b, c (Å)**<br>**α, β, γ (°)** | 101.34, 101.34, 101.34;<br>90.0, 90.0, 90.0 |
| **No. unique reflections** | 372727 (2386) |
| **Multiplicity** | 13.48 (14.13) |
| **Completeness (%)** | 100.0 (100.0) |
| **Mean I/sigma(I)** | 20.49 (1.31) |
| **Wilson B-factor (Å$^2$)** | 28.18 |
| **$R_{merge}$** | 0.068 (2.694) |
| **$R_{pim}$** | 0.019 (0.741) |
| **$CC_{1/2}$** | 1.000 (0.300) |
| **Refinement** | |
| **Resolution range (Å)** | 41.37–1.42 (1.46–1.42) |
| **No. reflections** | 32718 (2586) |
| **$R_{work}$** | 0.1735 (0.3057) |
| **$R_{free}$** | 0.1968 (0.3267) |
| **No. non-hydrogen atoms** | 1539 |
| **(poly)peptides** | 1282 |
| **ligands** | 42 |
| **solvent** | 215 |
| **RMSD bonds (Å)** | 0.006 |
| **RMSD angles (°)** | 0.856 |
| **Avg. B-factor (Å$^2$)** | 27 |
| **(poly)peptides** | 24 |
| **ligands** | 38 |
| **solvent** | 38 |
| **Ramachandran favored (%)** | 99.17 |
| **allowed (%)** | 0.83 |
| **outliers (%)** | 0.00 |
